# KAT6A acetylation drives metabolic adaptation to mediate cellular growth and mitochondrial metabolism through AMPK interaction

**DOI:** 10.1101/2025.01.14.633047

**Authors:** Mariko Aoyagi Keller, Andreas Ivessa, Tong Liu, Hong Li, Peter J Romanienko, Michinari Nakamura

## Abstract

Diets influence metabolism and disease susceptibility, with lysine acetyltransferases (KATs) serving as key regulators through acetyl-CoA. We have previously demonstrated that a ketogenic diet alleviates cardiac pathology, though the underlying mechanisms remain largely unknown. Here we show that KAT6A acetylation is crucial for mitochondrial function and cell growth. Proteomic analysis revealed that KAT6A is acetylated at lysine (K)816 in the hearts of mice fed a ketogenic diet under hypertension, which enhances its interaction with AMPK regulatory subunits. RNA-sequencing analysis demonstrated that the KAT6A acetylation-mimetic mutant stimulates AMPK signaling in cardiomyocytes. Moreover, the acetylation-mimetic mutant mitigated phenylephrine-induced mitochondrial dysfunction and cardiomyocyte hypertrophy via AMPK activation. However, KAT6A-K816R acetylation-resistant knock-in mice unexpectedly exhibited smaller hearts with enhanced AMPK activity, conferring protection against neurohumoral stress-induced cardiac hypertrophy and remodeling. These findings indicate that KAT6A regulates metabolism and cellular growth by interacting with and modulating AMPK activity through K816-acetylation in a cell type-specific manner.

## INTRODUCTION

Dietary nutrients are critical for maintaining organ structure and function by modulating cellular metabolite levels. Consumption of unhealthy diets disrupts metabolic homeostasis, characterized by impaired energy metabolism, dysregulated signaling pathways, lipotoxicity, and glucotoxicity^1, 2, 3^, thereby contributing to cardiometabolic diseases^4^, such as cardiac hypertrophy, atherosclerosis, and contractile dysfunction. Fatty acids serve as key energy substrates, major membrane components, and signaling molecules in the heart. Additionally, other diet-derived metabolites act as alternative energy sources, signaling molecules, or bioactive metabolites, influencing cardiovascular health and disease pathogenesis^5, 6, 7^. Our previous work demonstrated that elevating ketone body levels through a low-carbohydrate, high-fat ketogenic diet (KD) prevents high blood pressure-induced cardiac remodeling and heart failure in mice. This cardioprotective effect correlates with increased lysine acetylation in non-histone proteins from whole heart lysates, as well as in proteins localized to the cytosol, nuclei, and mitochondria^8^. These findings underscore the pivotal role of diet-mediated lysine acetylation, as observed with a KD, in promoting heart health.

Excessive lysine acetylation of mitochondrial proteins in the heart has been observed during early stages of heart failure in mouse models and in end-stage failing human hearts^9^. In line with these findings, increased mitochondrial protein acetylation caused by a reduced NAD^+^/NADH ratio in cardiac-specific Ndufs4 knockout (KO) mice has been shown to sensitize the mitochondrial permeability transition pore, thereby accelerating heart failure^10^. In contrast, a recent study using double KO mice for carnitine acetyltransferase and sirtuin 3 showed no association between mitochondrial hyperacetylation in the heart and susceptibility to heart failure^11^. Furthermore, modulating NAD^+^ levels was shown to alleviate heart failure independently of mitochondrial protein acetylation levels^12^, highlighting the complex and sometimes contradictory roles of lysine acetylation in heart failure. These discrepancies emphasize the need for further investigation into the effects of site-specific acetylation on individual protein function and its implications for cardiac function.

The thickness of the heart muscle is tightly regulated under physiological conditions to maintain adequate blood flow to peripheral organs^13^. However, prolonged hypertensive or dietary stress can lead to increased wall thickness of the heart, termed pathological hypertrophy, which is partially driven by dysregulated post-translational modifications^14, 15^. The variable effects of lysine acetylation on cardiac function led us to hypothesize that specific lysine acetylation events, rather than global protein acetylation, are critical for ketone body-mediated regulation of cardiac hypertrophy and function. To this end, we conducted an unbiased proteomic analysis using liquid chromatography-tandem mass spectrometry (LC-MS/MS). We identified acetylation of lysine acetyltransferase 6A (KAT6A) at lysine 816 (K816), located in exon 16, in the hearts of mice fed a KD under high blood pressure, but not in those on normal chow (NC). KAT6A catalyzes the transfer of acetyl groups from acetyl-CoA to lysine residues^16^. Pathogenic variants in the *KAT6A* gene cause KAT6A syndrome, featuring neurodevelopmental disorders, hypotonia, and heart abnormalities^17^. Notably, while dysfunction of the KAT6A catalytic subunit severely impacts transcriptional regulation, genotype-phenotype correlations reveal that late-truncating pathogenic variants (in exon 16-17, encoding the non-catalytic acidic domain) are significantly more prevalent^17^. In this study, we sought to elucidate the functional role and significance of KAT6A-K816 acetylation in mitochondrial and cardiac function, hypothesizing that it serves as a central hub for ketone body-mediated regulation of cardiac function.

## RESULTS

### Ketone body increases acetylation of KAT6A at K816 in the hearts of hypertensive mice

Since a KD specifically impacts diseased hearts, such as those with hypertensive heart disease^8^, we focused on this pathological condition. To identify proteins undergoing acetylation in the hearts of C57BL/6J wild-type (WT) mice fed a KD compared to a control NC diet under a high blood pressure condition, we performed LC-MS/MS proteomic analysis. We identified 966 acetylated peptides corresponding to 320 acetylated proteins in the hearts of mice fed an NC diet (Extended Data Table 1), while 1597 acetylated peptides corresponding to 425 acetylated proteins were identified in the hearts of mice fed a KD (Extended Data Table 2). Consistent with previously reported immunoblot data^8^, a KD diet increased the overall number of acetylated proteins and lysine residues in the mouse heart (Fig. 1a). Lysine acetylation is regulated by lysine acetyltransferases (KATs) and deacetylases, depending on intracellular levels of acetyl-CoA and NAD^+16^. Gene ontology (GO) molecular function enrichment analysis of 170 acetylated proteins observed in mice fed a KD revealed significant enrichment in GO terms related to acetyltransferase activity, pyridine nucleotide redox regulation, and metabolic processes (Fig. 1b).

**Fig. 1:**
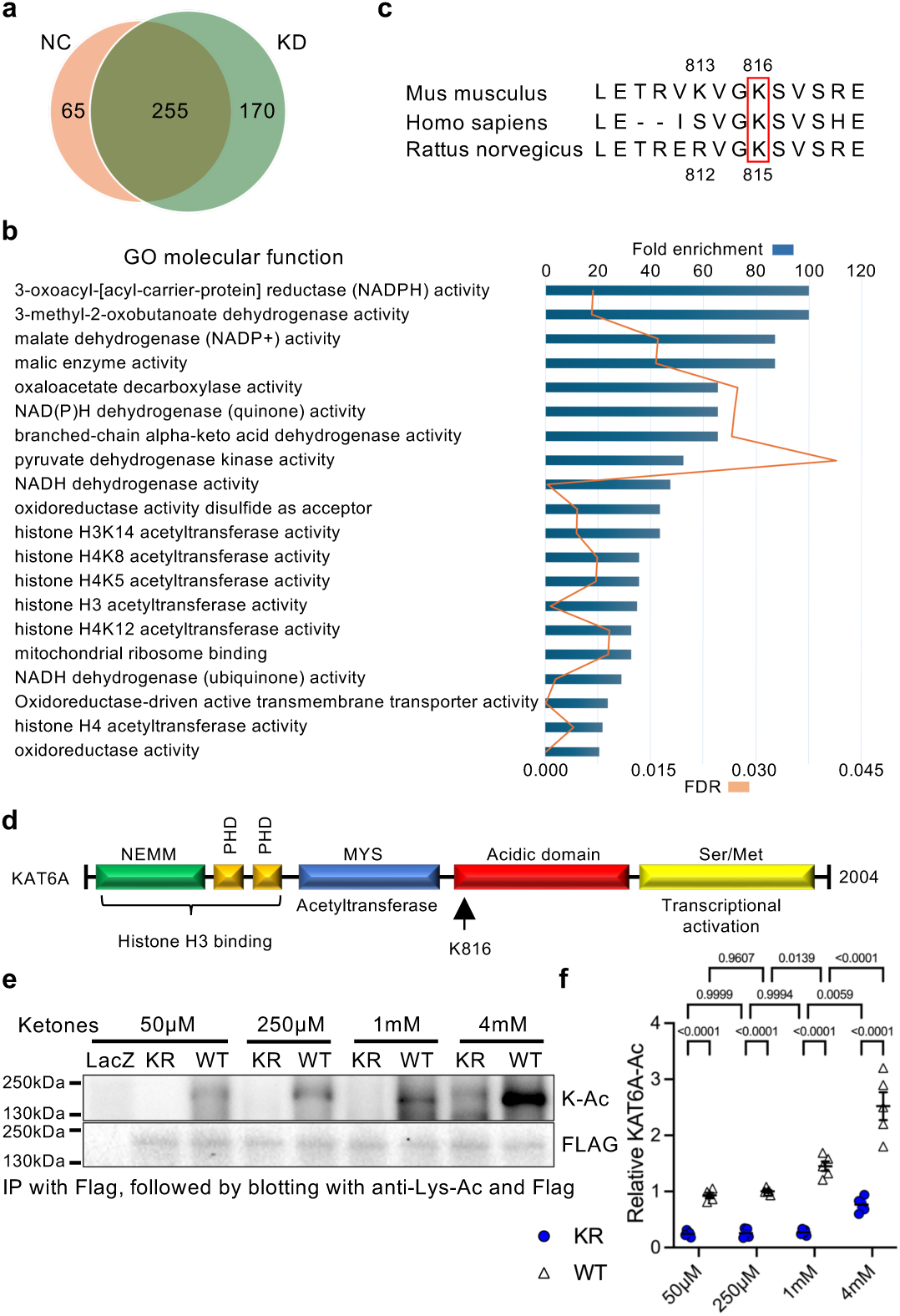
Lysine acetyltransferase 6A (KAT6A) is acetylated at K816 in the hearts of mice fed a ketogenic diet under conditions of high blood pressure. **a,** Venn diagram showing the number of acetylated proteins in the hearts of mice fed a ketogenic diet (KD) or normal chow (NC) under conditions of high blood pressure. **b**, Gene ontology (GO) molecular function enrichment analysis using 170 acetylated proteins uniquely identified in mice on a KD diet. **c**, Amino acid sequences highlighting the conservation of the KAT6A K816 acetylation site across various species. **d,** Schematic illustration of the KAT6A amino acid domain structure. **e-f**, Immunoblots showing KAT6A-K815 acetylation in response to ketone bodies (a cocktail of β-hydroxybutyrate and acetoacetate) at the indicated concentrations in cardiomyocytes (**e**). Rat neonatal cardiomyocytes were transfected with adenovirus harboring FLAG-tagged human KAT6A-wild type (WT) or K815R (KR) for two days, followed by treatment with ketones for four hours. Immunoprecipitation was performed using FLAG agarose beads, followed by immunoblots with anti-Lys-Ac and FLAG antibodies. Densitometric analysis of the immunoblots (n=5) (**f**). Two-way ANOVA followed by Tukey’s multiple comparison test. *N* represents biologically independent replicates. *p* values are shown in the figure. Data are mean ± SEM. Source data are provided as a Source Data file.

To further investigate the mechanism by which a KD enhances acetylation in the heart, we examined the acetylation status of lysine acetyltransferases (KATs). Among the MYST, GNAT, and p300/CBP families of canonical KATs^16^, KD-mediated lysine acetylation in the heart was detected in KAT6A and EP300 (Extended Data Table 3). LC-MS/MS mass-spectrometry analysis identified endogenous KAT6A acetylation at K813 (located in exon 15) and K816 (located in exon 16) in the hearts of mice fed a KD (Extended Data Fig. 1a). Given the clinical relevance and high prevalence of mutations in exons 16-17 of KAT6A, which encompass the non-catalytic acidic domain, we aimed to elucidate the role of KAT6A-K816 acetylation in cardiac disease. The K816 of KAT6A is highly conserved across species: in *Mus musculus*, it corresponds to K815 in *Homo sapiens* and *Rattus norvegicus*. In contrast, KAT6A-K813 in *Mus musculus* is not conserved in other species (Fig. 1c). K816 is located within the acidic domain, a region that may influence protein-protein interactions (Fig. 1d).

To establish the link between KAT6A-K816 acetylation and ketone bodies, while excluding other potential factors related to a KD, we conducted *in vitro* experiments using adenovirus (Ad)-mediated expression of human KAT6A-wild type (WT), K815R (acetylation-resistant, corresponding to mouse K816), and K815Q (acetylation-mimicking) mutants in primary rat neonatal cardiomyocytes. We have previously demonstrated that ketone bodies increase lysine acetylation levels in whole protein lysates as well as in subcellular fractions, including cytosolic, mitochondrial, and nuclear proteins, in a dose-dependent manner^8^. Here, we first examined whether ketone bodies alter the acetylation status of KAT6A at K815 in cardiomyocytes. Treatment of cardiomyocytes with ketone bodies (β-hydroxybutyrate and acetoacetate) resulted in a dose-dependent increase in KAT6A-K815 acetylation (Fig. 1e,f), indicating that the KD-mediated increase in KAT6A-K816 acetylation in the heart is induced by elevated ketone body levels. Therefore, we sought to elucidate the roles of ketone body-mediated KAT6A-K815 acetylation in cardiac physiology and pathology, particularly in cardiac hypertrophy and contractile dysfunction.

### Acetylation of KAT6A at K815 stimulates AMPK signaling

Next, we investigated whether KAT6A-K815 acetylation influences histone and non-histone acetylation. Overexpression of neither KAT6A-WT nor K815 mutants altered overall lysine acetylation levels (Extended Data Fig. 1b); however, acetylation of histone 3 (H3)K9 and H3K14 was increased by KAT6A overexpression, with no overt differences between WT and K815 mutants (Extended Data Fig. 1c). These findings suggest that KAT6A specifically promotes H3K9 and H3K14 acetylation in cardiomyocytes independently of KAT6A-K815 acetylation. In line with these findings, KAT6A-WT and its mutants are localized in the nucleus in cardiomyocytes, irrespective of K815 acetylation status (Extended Data Fig. 1d).

Then, we investigated the functional role of KAT6A-K815 acetylation in cardiomyocytes. To unbiasedly understand which biological pathways are significantly activated or inhibited by KAT6A-K815 acetylation in cardiomyocytes, we conducted RNA-sequencing analysis. The heatmap shows significant alterations in gene expression by KAT6A-WT compared to control LacZ in cardiomyocytes (Fig. 2a). The gene set enrichment analysis (GSEA) using the C2.All.v6.2 (curated) gene sets database revealed enrichment of gene sets related to the activation of fatty acid oxidation by AMP-activated protein kinase (AMPK) (REACTOME) and the regulation of autophagy (KEGG) (Fig. 2b and Extended Data Table 4). Furthermore, the effect of K815 acetylation was evaluated by expressing KAT6A-K815Q or K815R mutants (Fig. 2c). The GSEA showed that gene sets associated with the regulation of mTOR by AMPK and AMPK-stimulated catabolic processes are significantly enriched in cardiomyocytes expressing KAT6A-K815Q compared to those expressing KAT6A-K815R (Fig. 2d and Extended Data Table 5). In line with these results, KAT6A-K815Q expression was sufficient to significantly increase phosphorylation of Acetyl-CoA carboxylase (ACC) at Ser79, a well-known AMPK substrate, compared to LacZ or KAT6A-K815R (Fig. 2e). In addition, ketone body-mediated activation of AMPK was suppressed in cardiomyocytes expressing the KAT6A-K815R mutant compared to those expressing KAT6A-WT (Fig. 2f). Notably, the concentrations of ketone bodies used in this experiment align with physiological and pathological conditions: 100-250 μM under normal conditions, 1-2 mM after 24 hours of fasting or prolonged exercise, 1-5 mM in mice and humans fed a ketogenic diet, and up to 20 mM in diabetic ketoacidosis. These findings indicate that ketone bodies increase acetylation of KAT6A at K815, which, in turn, enhances AMPK signaling in cardiomyocytes.

**Fig. 2:**
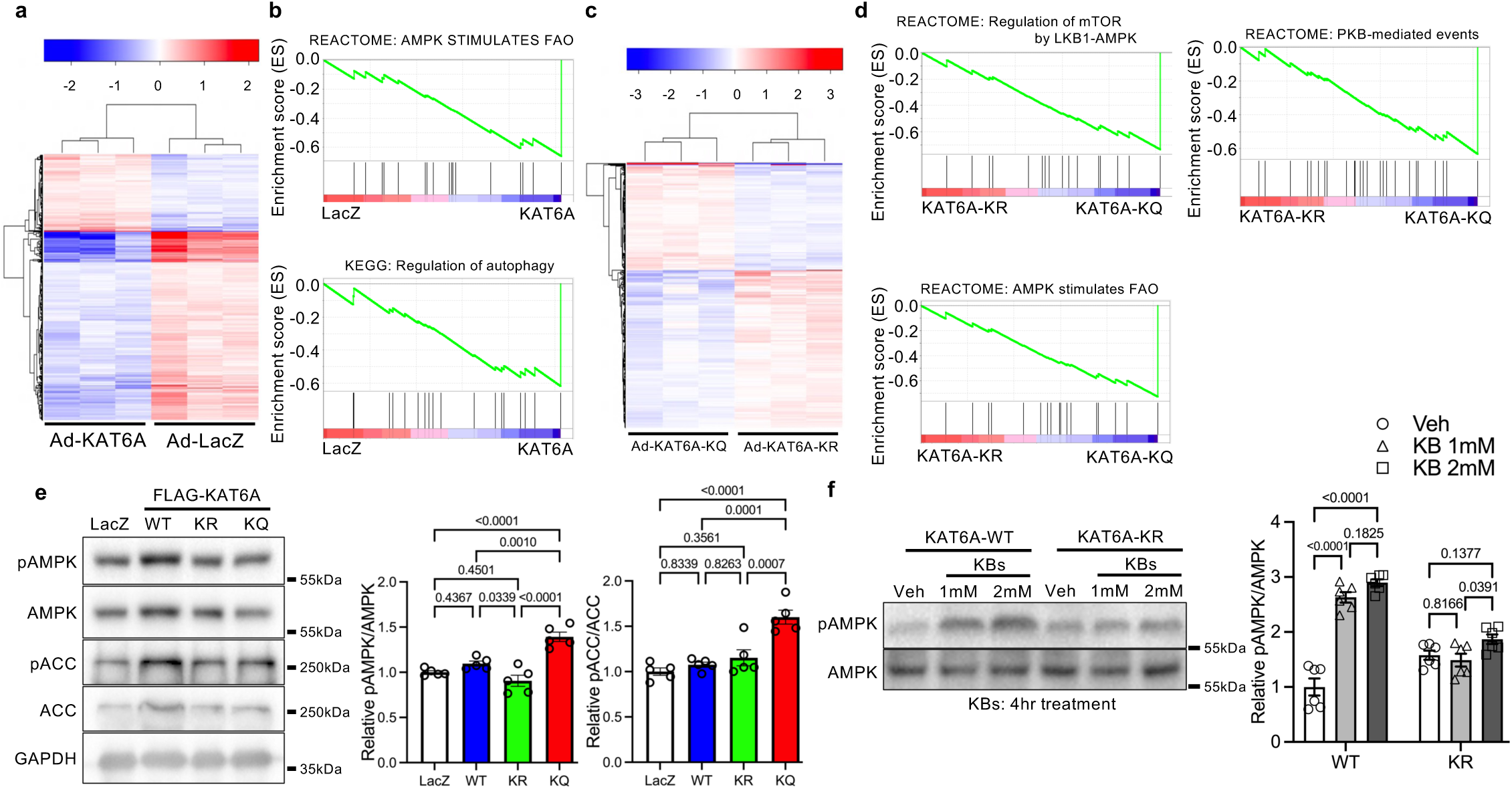
Acetylation of KAT6A enhances AMPK activity in cardiomyocytes. **a**, Heatmap of the top 500 differentially expressed genes in cardiomyocytes expressing KAT6A-wild type (WT) compared to LacZ as a control (n=3). **b**, Gene set enrichment analysis (GSEA) plots of the ‘AMPK stimulates fatty acid oxidation (FAO)’ (REACTOME) and ‘Regulation of autophagy’ (KEGG) pathways enriched in cardiomyocytes expressing KAT6A-WT compared to LacZ (see Extended Data Table 4). **c**, Heatmap of the top 500 differentially expressed genes in cardiomyocytes expressing KAT6A-K815Q (KQ) compared to KAT6A-K815R (KR) (n=3). **d**, GSEA plots of ‘Regulation of mTOR by LKB1-AMPK’ (REACTOME), ‘AMPK stimulates FAO’ (REACTOME), and ‘PKB-mediated events’ (REACTOME) pathways enriched in cardiomyocytes expressing KAT6A-K815Q (KQ) compared to KAT6A-K815R (KR) (see Extended Data Table 5). **e**, Immunoblots showing AMPK activation in cardiomyocytes (left) and their densitometric analysis (right) (n=5). **f**, Immunoblots showing AMPK activation in cardiomyocytes treated with ketone bodies (KB) at the indicated concentrations or vehicle for four hours (left) along with their densitometric analysis (right) (n= 6). One-way ANOVA followed by Tukey’s multiple comparison test. *N* represents biologically independent replicates. *p* values are shown in the figure. Data are mean ± SEM. Source data are provided as a Source Data file.

### KAT6A-K815 acetylation increases the interaction with AMPK subunits

Next, we investigated the underlying mechanisms by which KAT6A-K815 acetylation enhances AMPK activity. To identify changes in KAT6A interaction partners by K815 acetylation, we transfected Ad-FLAG-KAT6A-WT, -K815R, and -K815Q mutants into cardiomyocytes for 2 days. Ad-FLAG was used as a control. KAT6A was immunoprecipitated with α-FLAG affinity beads, followed by washing, competitive elution with 3x FLAG peptides, and separation by SDS-PAGE. LC-MS/MS proteomic analyses were performed to unbiasedly identify proteins interacting with KAT6A-WT and its mutants (Extended Data Table 6). To our surprise, and consistent with RNA-sequencing findings, mass spectrometry identified AMPK β/γ subunits as major interacting proteins with KAT6A in cardiomyocytes (Table 1). Notably, KAT6A-WT and the K815Q mutant interacted with AMPK regulatory subunits β1 and/or γ2 more efficiently than the K815R mutant. These findings suggest that KAT6A-K815 acetylation stimulates AMPK signaling through increased interactions between KAT6A and AMPK β/γ regulatory subunits.

**Table 1:** Top 5 proteins identified by mass-spectrometry analyses using the immunoprecipitation samples with flag agarose beads in cardiomyocytes expressing KAT6A-wild type (WT), K815R (KR), or K815Q (KQ) mutants.

| KR top 5 |  | WT top 5 |  | KQ top 5 |  |
| --- | --- | --- | --- | --- | --- |
| Protein | Ratio (KR/Ctrl) | Protein | Ratio (WT/Ctrl) | Protein | Ratio (KQ/Ctrl) |
| KAT6A | 14 | KAT6A | 25.5 | AMPKβ1 | 25.5 |
| AMPKγ1 | 5.2 | AMPKβ1 | 18.5 | AMPKγ1 | 19 |
| KARS | 5 | AMPKγ1 | 12.4 | IMPORTIN7 | 11 |
| CCT8 | 4.5 | NARDILYSIN | 9.5 | UBE2O | 11 |
| D3ZJE2 | 4.5 | IMPORTIN7 | 8 | AMPKγ2 | 9.4 |

### KAT6A-K815 acetylation enhances mitochondrial respiratory capacity and fatty acid oxidation through AMPK in cardiomyocytes

AMPK plays a key role in maintaining cardiac function and morphology by regulating metabolism and mitochondrial activity. To explore the functional significance of KAT6A-K815 acetylation in cardiomyocyte size, mitochondrial function, and substrate metabolism, we transduced rat neonatal cardiomyocytes with adenoviruses harboring YFP-tagged KAT6A-K815R or YFP-tagged KAT6A-K815Q mutants. We utilized YFP-tagged KAT6A to identify cardiomyocytes expressing mutant KAT6A. At baseline, no significant differences in size were observed between cardiomyocytes expressing the KAT6A-K815R and KAT6A-K815Q mutants (Fig. 3a). However, despite the absence of changes in cardiomyocyte size, mitochondrial energy metabolism was markedly altered in cardiomyocytes expressing KAT6A-K815 mutants under non-hypertrophic conditions (Fig. 3b). Seahorse experiments showed that the basal oxygen consumption rate (OCR), ATP-linked respiration, and maximal respiratory capacity are significantly elevated in H9C2 cardiomyoblast cells expressing the KAT6A-K815Q mutant compared to those expressing the KTA6A-K815R mutant (Fig. 3bc). These effects were diminished in the KAT6A-K815Q group but not in the KAT6A-K815R group following adenovirus-mediated overexpression of dominant-negative AMPK (Fig. 3c), indicating that AMPK functions downstream of KAT6A-K815 acetylation.

**Fig. 3:**
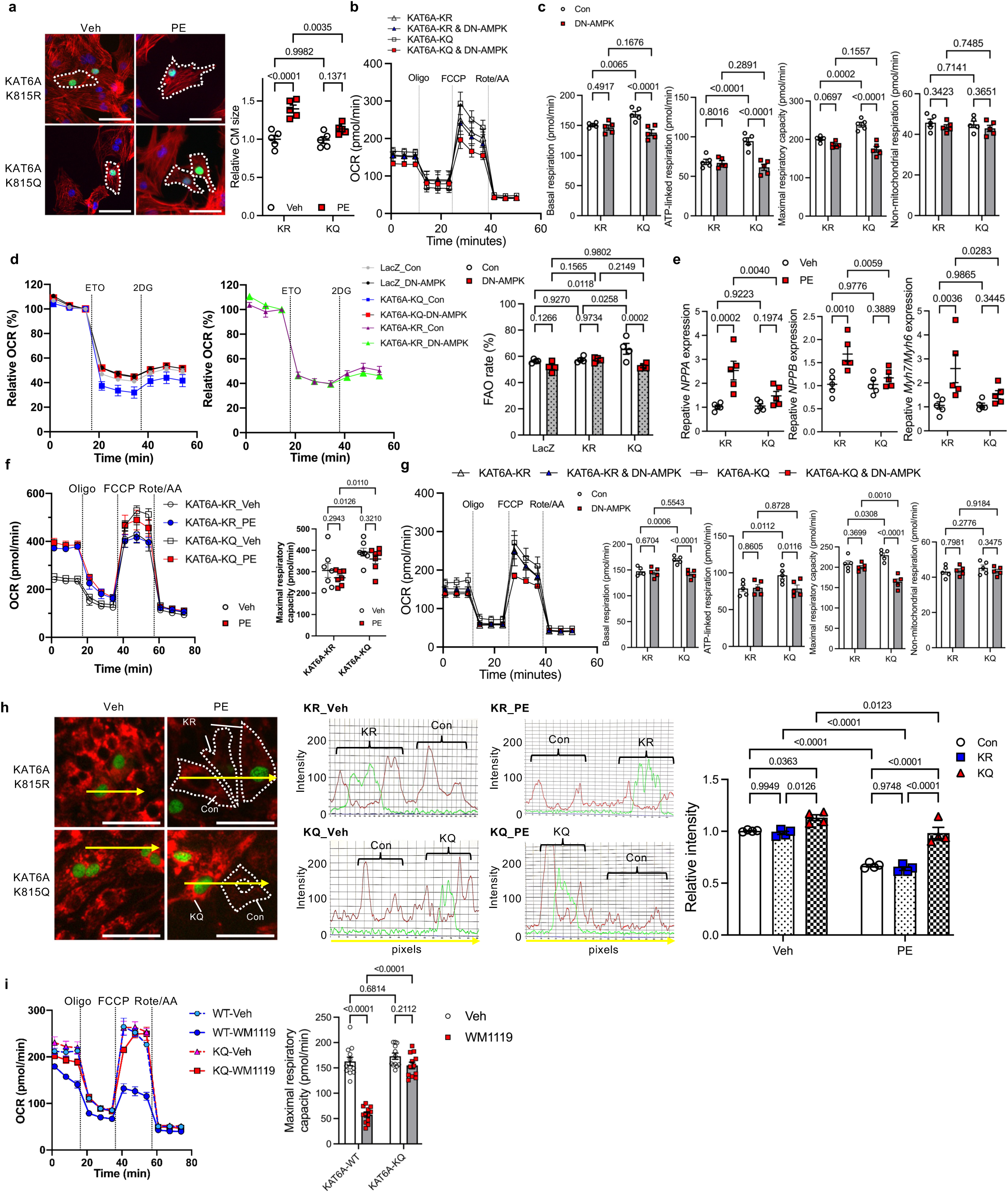
Acetylation of KAT6A suppresses hypertrophy and mitochondrial dysfunction with enhancing fatty acid oxidation in an AMPK-dependent manner in cardiomyocytes. **a**, Representative images of cardiomyocytes (CMs) expressing YFP-tagged KAT6A-K815R (KR) or - K815Q (KQ) in response to phenylephrine (PE) or vehicle (Veh) treatment for 48 hours. Dashed lines outline individual CMs transfected with adenovirus. Scale bar, 50 μm. Relative CM size (right) (n=5). **b-c**, Mitochondrial function assessed by Seahorse experiments. Representative oxygen consumption rate (OCR) plot in CMs transfected with KAT6A-K815R or -K815Q, co-expressed with either dominant-negative AMPK (DN-AMPK) or LacZ as a control (b). Histograms showing basal respiration, ATP-linked respiration, maximal respiratory capacity, and non-mitochondrial respiration (n=5) (c). **d**, Representative OCR plot evaluating fatty acid oxidation (FAO) rates in CMs transfected with KAT6A-K815R, KAT6A-K815Q, or LacZ as a control, in the presence of DN-AMPK or LacZ. Histograms showing etomoxir (ETO)-inhibitable FAO rates (n=4). **e**, Relative gene expression of *NPPA*, *NPPB*, and *MYH7/MYH6* ratio in CMs expressing KAT6A-K815R or -K815Q in response to PE or vehicle for 48 hours (n=5). **f**, Seahorse experiments showing mitochondrial function in response to PE for 24 hours. Representative OCR plot of CMs transfected with KAT6A-K815R or -K815Q. Histograms showing maximal respiratory capacity (n=7). **g**, Seahorse experiments showing mitochondrial function under PE treatment for 24 hours. Representative OCR plot of CMs transfected with KAT6A-K815R or -K815Q in the presence of DN-AMPK or LacZ (left). Histograms showing basal respiration, ATP-linked respiration, maximal respiratory capacity, and non-mitochondrial respiration (right) (n=5). **h**, Tetramethylrhodamine methyl ester (TMRM) images, signal intensity of TMRM and YFP, and histograms quantifying relative TMRM intensity in CMs expressing YFP-KAT6A-K815R or -K815Q under PE or vehicle treatment for 24 hours (n=4). Scale bar, 50 μm. **i**, Representative OCR plot of CMs transfected with KAT6A-WT or -K815Q in the presence of WM1119 or vehicle under PE treatment for 24 hours. Histograms showing maximal respiratory capacity (n=12). Two-way ANOVA followed by Tukey’s multiple comparison test. *N* represents biologically independent replicates. *p* values are shown in the figure. Data are mean ± SEM. Source data are provided as a Source Data file.

AMPK is known to promote fatty acid β-oxidation by reducing the production of malonyl-CoA, an endogenous inhibitor of mitochondrial carnitine palmitoyl transferase 1 (CPT1), via phosphorylation of acetyl-CoA carboxylase 1 (ACC1) at Serine 79. Consistent with RNA-sequencing results, fatty acid oxidation, as assessed by etomoxir (ETO)-inhibitable OCR in Seahorse experiments, was significantly increased in cardiomyocytes expressing the KAT6A-K815Q mutant but not in those expressing the KAT6A-K815R mutant. These effects were also reversed by adenovirus-mediated inhibition of AMPK activity (Fig. 3d). Taken together, these findings suggest that KAT6A-K815 acetylation enhances mitochondrial function and fatty acid oxidation through AMPK activation in cardiomyocytes.

### KAT6A-K815 acetylation attenuates phenylephrine-induced hypertrophy and mitochondrial dysfunction in cardiomyocytes

Next, to test the hypothesis that acetylation of KAT6A at K815 is involved in cell growth and mitochondrial function in response to hypertrophic stimuli, cardiomyocytes were treated with phenylephrine, an α-adrenergic receptor agonist that mimics hypertensive conditions. Forty-eight-hour phenylephrine treatment significantly increased cell size in cardiomyocytes expressing the YFP-KAT6A-K815R mutant, while cardiomyocytes expressing the YFP-KAT6A-K815Q mutant exhibited a markedly attenuated hypertrophic response (Fig. 3a). Consistent with these observations, the phenylephrine-induced fetal gene expression, including *Nppa*, *Nppb*, and the ratio of *Myh7*/*Myh6*, was suppressed in cardiomyocytes expressing the KAT6A-K815Q mutant compared to those expressing the KAT6A-K815R mutant (Fig. 3e). These findings suggest that KAT6A-K815 acetylation mitigates hypertrophic responses in cardiomyocytes under stress conditions.

Phenylephrine treatment for 24 hours increased basal OCR in cardiomyocytes expressing either the KAT6A-K815Q or KAT6A-K815R mutant, while phenylephrine-induced mitochondrial dysfunction, characterized by a reduction in maximal respiratory capacity, was significantly attenuated in cardiomyocytes expressing the KAT6A-K815Q mutant compared to those expressing the KAT6A-K815R mutant (Fig. 3f). The protective effect of the KAT6A-K815Q mutant on mitochondrial function under phenylephrine treatment was abolished by adenovirus-mediated overexpression of dominant-negative AMPK (Fig. 3g). To further assess mitochondrial function, mitochondrial membrane potential was measured using Tetramethylrhodamine methyl ester (TMRM) in rat neonatal cardiomyocytes expressing either the YFP-KAT6A-K815Q or YFP-KAT6A-K815R mutant, which identify cells expressing a mutant protein. In line with the Seahorse data, cardiomyocytes expressing the YFP-KAT6A-K815R mutant exhibited a significant reduction in mitochondrial membrane potential in response to phenylephrine, as indicated by a lower intensity of TMRM signal. In contrast, cardiomyocytes expressing the YFP-KAT6A-K815Q mutant maintained mitochondrial membrane potential even in the presence of phenylephrine (Fig. 3h). These findings indicate that KAT6A-K815 acetylation mitigates phenylephrine-induced mitochondrial dysfunction by enhancing AMPK activity.

Finally, we asked how the catalytic activity of KAT6A contributes to mitochondrial function through KAT6A-K815 acetylation in cardiomyocytes. We hypothesized that KAT6A-K815 acetylation enhances energy homeostasis independently of its MYST catalytic activity, given the minimal impact of KAT6A-K815 acetylation on lysine acetylation (Extended Data Fig. 1bc). To test this, cardiomyocytes were transfected with Ad-KAT6A-WT or Ad-KAT6A-K815Q in the presence or absence of WM-1119, an inhibitor of KAT6A catalytic activity^18^, followed by Seahorse experiments. As reported^18^, WM-1119 dose-dependently reduced H3K14 acetylation without influencing H3K9 acetylation (Extended Data Fig. 1e). We found that WM-1119 markedly reduces maximal respiration in cardiomyocytes expressing KAT6A-WT, suggesting that the MYST catalytic activity of KAT6A is necessary for maintaining energy homeostasis in cardiomyocytes (Fig. 3i). Importantly, however, WM-1119 failed to impair maximal respiration in cardiomyocytes expressing the KAT6A-K816Q mutant. These results indicate that the catalytic activity of KAT6A is dispensable for enhancing maximal respiration in cardiomyocytes once KAT6A is acetylated at K815, potentially due to auto-acetylation of KAT6A at this site. Taken together, these findings suggest that while KAT6A catalytic activity is essential for the maintenance of mitochondrial function and stress resistance in cardiomyocytes, KAT6A-K815 acetylation provides a catalytic-independent mechanism for enhancing mitochondrial function. This mechanism likely involves the promotion of KAT6A’s scaffolding role in AMPK complex formation and activation through K815 acetylation.

### KAT6A-K816 acetylation-resistant knock-in mice display the smaller heart size with preserved cardiac function

To investigate the functional significance of KAT6A-K815 acetylation *in vivo*, we generated KAT6A-K816R (corresponding to human KAT6A-K815) acetylation-resistant knock-in mice using the CRISPR-CAS9 system at the Rutgers Genome Editing Shared Resource. Successful generation of the knock-in mice was confirmed by next-generation sequencing (Extended Data Fig. 2a). The KAT6A-K816R knock-in mice were backcrossed with C57BL/6J WT mice for more than six generations. The KAT6A-K816R homozygous knock-in mice were born at Mendelian ratios and exhibited normal viability (Extended Data Table 7), suggesting that acetylation of KAT6A at K816 is dispensable for normal embryonic development in mice. Consequently, homozygous knock-in mice were used for all subsequent experiments.

To characterize the newly generated mouse line, we first evaluated systemic phenotypes, including body weight, organ weights, and blood levels of glucose and β-hydroxybutyrate, in both sexes. At 12 weeks of age, there were no significant differences in body weight between KAT6A-K816R homozygous knock-in mice and their littermate WT controls, either at baseline or after fasting, in both sexes (Fig. 4a,b). Fasting blood sugar levels were also comparable between the two groups in both sexes (Fig. 4c). Although KAT6A-K816 acetylation increased in response to a ketogenic diet, inhibition of KAT6A K816 acetylation did not affect serum β-hydroxybutyrate levels in either sex at 12 weeks of age (Fig. 4d).

**Fig. 4:**
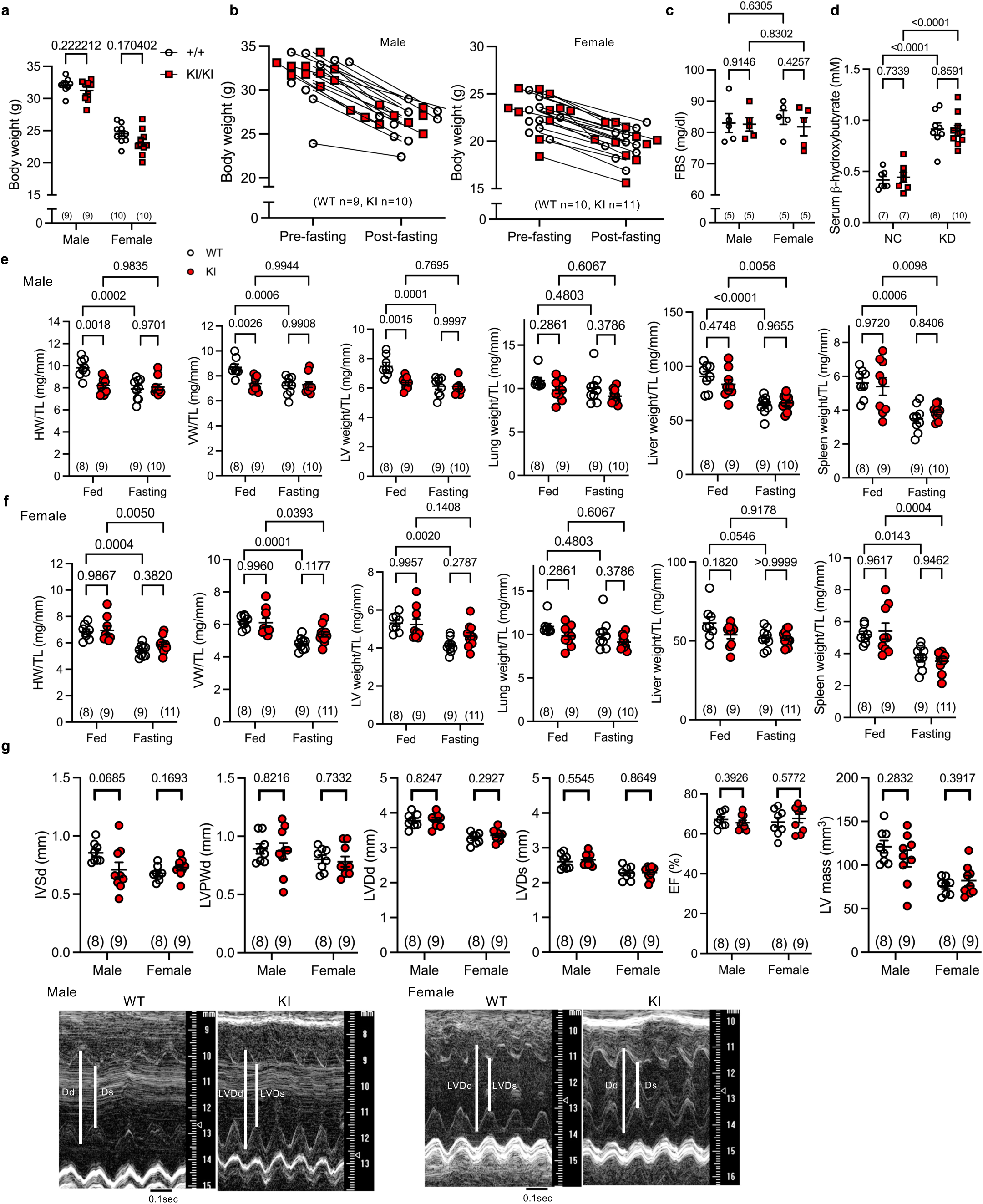
KAT6A-K816R acetylation-resistant knock-in mice exhibit smaller hearts in young males, but not females, with preserved cardiac function. **a**, Body weight of male and female wild-type (WT, +/+) and knock-in (KI, KI/KI) mice at 12 weeks of age. Unpaired *t* test within each sex. **b**, Body weight changes following a 36-hour fasting in male and female mice at 12 weeks of age. **c-d**, Fasting blood sugar levels (c) and serum β-hydroxybutyrate levels (d) in the indicated mice fed either normal chow (NC) or a ketogenic diet (KD). **e-f**, Organ weights normalized to tibia length (TL) in KI and WT littermate mice of both sexes at 8 weeks of age (e: males; f: females). Weights measured include heart weight (HW), ventricular weight (VW), left ventricular (LV) weight, lung weight, liver weight, and spleen weight. Two-way ANOVA followed by Tukey’s multiple comparison test (c-f). **g**, Echocardiographic measurements, including interventricular septal wall thickness at end-diastole (IVSd), LV posterior wall thickness at end-diastole (LVPWd), LV end-diastolic diameter (LVDd), LV end-systolic diameter (LVDs), ejection fraction (EF), and LV mass, in KI and WT littermate mice of both sexes at 8 weeks of age. Unpaired *t* test within each sex. Representative M-mode echocardiography images are shown in the lower panel. The number in brackets indicates the sample number. *N* represents biologically independent replicates. *p* values are shown in the figure. Data are mean ± SEM. Source data are provided as a Source Data file.

Then, we evaluated various organ weights normalized to tibia length in both sexes at 8 weeks of age. Surprisingly, heart weight, ventricular weight, and left ventricular weight were significantly smaller in male KAT6A-K816R knock-in mice compared to their littermate WT controls under fed conditions (Fig. 4e). This unexpected reduction in heart size was not observed after 36 hours of fasting or in female KAT6A-K816R knock-in mice (Fig. 4e,f). Additionally, the weights of other organs, including the lungs, liver, and spleen, were unaffected by the inhibition of KAT6A-K816 acetylation in either sex (Fig. 4e,f). Consistent with the reduced heart weight in fed male knock-in mice, interventricular septal wall thickness at diastole was thinner in KAT6A-K816R knock-in male, as assessed by echocardiography, whereas no such difference was observed in females (Fig. 4g). Despite the smaller heart size, cardiac function remained comparable between KAT6A-K816R knock-in and WT mice of both sexes at 8 weeks of age (Fig. 4g). These findings indicate that KAT6A-K816 acetylation is essential for maintaining heart size in male mice at baseline *in vivo*, despite its lack of impact on cardiomyocyte size under non-stress condition *in vitro*. However, KAT6A-K816 acetylation appears dispensable for preserving baseline cardiac function in both sexes *in vivo*.

### KAT6A-K816 acetylation-resistant knock-in mice show increased AMPK signaling in the heart with smaller cardiomyocytes

We further examined mutant hearts under fasting and non-fasting conditions at 8 weeks of age using histological analyses. Consistent with autopsy and echocardiography data (Fig. 4), individual cardiomyocyte size was significantly smaller in the hearts of male KAT6A-K816R knock-in mice compared to their littermate WT controls under fed conditions. However, this difference was not observed in female mice (Fig. 5a,b). Notably, these size differences were undetectable under fasting conditions in both sexes. In contrast, no significant differences in cardiac fibrosis were observed between KAT6A-K816R knock-in and littermate WT controls, regardless of sex or fasting condition (Fig. 5c,d).

**Fig. 5:**
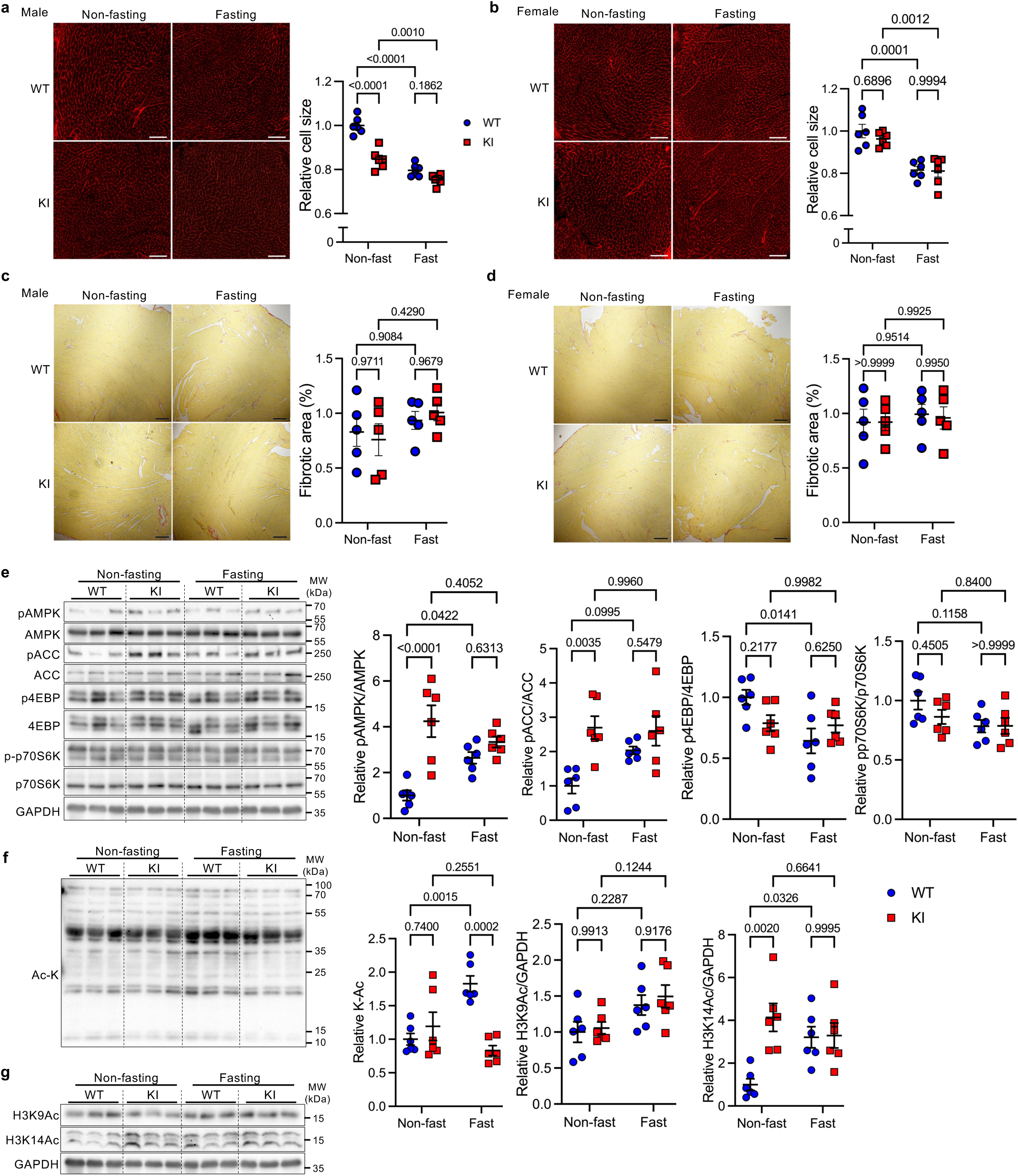
KAT6A-K816R acetylation-resistant knock-in mice exhibit smaller cardiomyocytes in young males, but not females, with increased AMPK activity. **a-b**, Wheat germ agglutinin (WGA) staining of heart sections from male (a) and female (b) homozygous KI and littermate WT mice under non-fasting and fasting conditions at 8 weeks of age. Quantitative analysis of relative cardiomyocyte size (n=6). Scale bar, 100 μm. **c**-**d**, Picric Acid Sirius Red (PASR) staining of heart sections from male (c) and female (d) homozygous KI and littermate WT mice under non-fasting and fasting conditions at 8 weeks of age. Quantitative analysis of fibrotic areas (n=5). Scale bar, 200 μm. **e**-**g**, Immunoblots showing AMPK and mTOR signaling pathways (e), overall lysine acetylation (K-Ac) (f), and histone H3K9 and H3K14 acetylation (g) in the hearts of homozygous KI and littermate WT male mice under non-fasting and fasting conditions at 8 weeks of age. Histograms showing their densitometric analyses (right, n=6). Two-way ANOVA followed by Tukey’s multiple comparison test. *N* represents biologically independent replicates. *p* values are shown in the figure. Data are mean ± SEM. Source data are provided as a Source Data file.

To investigate the mechanisms underlying the discrepancy between the *in vitro* phenotypes of cardiomyocytes and *in vivo* phenotypes of mutant hearts, we evaluated AMPK signaling and lysine acetylation status in the hearts of KAT6A-K816R knock-in mice by immunoblots. Under fed conditions, phosphorylation of AMPK and its substrate ACC phosphorylation was elevated in the hearts of KAT6A-K816R knock-in mice, but these differences were diminished after 36-hour fasting (Fig. 5e). In line with these observations, phosphorylation of 4EBP and p70S6K, downstream targets of mTOR, was slightly reduced in these hearts. These findings indicate that inhibition of KAT6A-K816 acetylation enhances AMPK activity, mimicking the fasting phenotype, which contrasts with the findings observed in cardiomyocytes *in vitro*. However, interestingly, the fasting-induced increase in overall lysine acetylation was suppressed in the hearts of KAT6A-K816R knock-in mice (Fig. 5f). While H3K9 acetylation levels were unaffected by fasting or the knock-in mutation (Fig. 5g), H3K14 acetylation levels were elevated in the hearts of KAT6A-K816R knock-in mice under fed conditions, resembling the increase seen during fasting. These findings indicate that inhibition of KAT6A-K816 acetylation *in vivo* alters AMPK signaling and lysine acetylation patterns, potentially contributing to the observed phenotypic differences between *in vitro* and *in vivo* models.

### Inhibition of KAT6A-K816 acetylation mitigates cardiac hypertrophy and dysfunction in response to hypertrophic stimuli in mice in vivo

To demonstrate the significance of KAT6A-K816 acetylation in the setting of chronic hypertension *in vivo*, we continuously administered neurohumoral stimulation (angiotensin II and phenylephrine) to KAT6A-K816R knock-in and age-matched control WT mice via osmotic pump for 3 weeks (Fig. 6a). Given that heart and ventricular weights are slightly but significantly smaller in KAT6A-K816 knock-in male mice compared to WT mice at 8 weeks of age, but the differences get smaller after the age, 10-week-old male knock-in and age-matched control WT mice were used in this experiment. There was no significant difference in heart weight and other organ weights, including lung, liver, and spleen, between both groups after 3 weeks of treatment with vehicle (Fig. 6b). While chronic neurohumoral stimulation induced cardiac hypertrophy in control WT mice, KAT6A-K816R knock-in mice suppressed the development of hypertrophy (Fig. 6b). Consistent with autopsy data, angiotensin II/phenylephrine-induced increases in interventricular and left ventricle (LV) posterior wall thickness at diastole and LV mass, as assessed by echocardiography, were suppressed by inhibition of KAT6A-K816 acetylation (Fig. 6c). Additionally, KAT6A-K816R knock-in mice attenuated neurohumoral stimulation-induced cardiac remodeling and systolic dysfunction (Fig. 6c). These findings were supported by histological analyses. While individual cardiomyocyte size was similar in response to vehicle between both groups, cardiomyocyte hypertrophy was attenuated in KAT6A-K816R knock-in mouse hearts compared to WT mice (Fig. 6d). Furthermore, inhibition of KAT6A-K816 acetylation mitigated cardiac fibrosis in response to angiotensin II and phenylephrine (Fig. 6e). These results suggest that acetylation of KAT6A at K816 promotes pathological cardiac hypertrophy, remodeling, and cardiac dysfunction in response to chronic hypertension, which is opposite to the findings observed in cardiomyocytes *in vitro*.

**Fig. 6:**
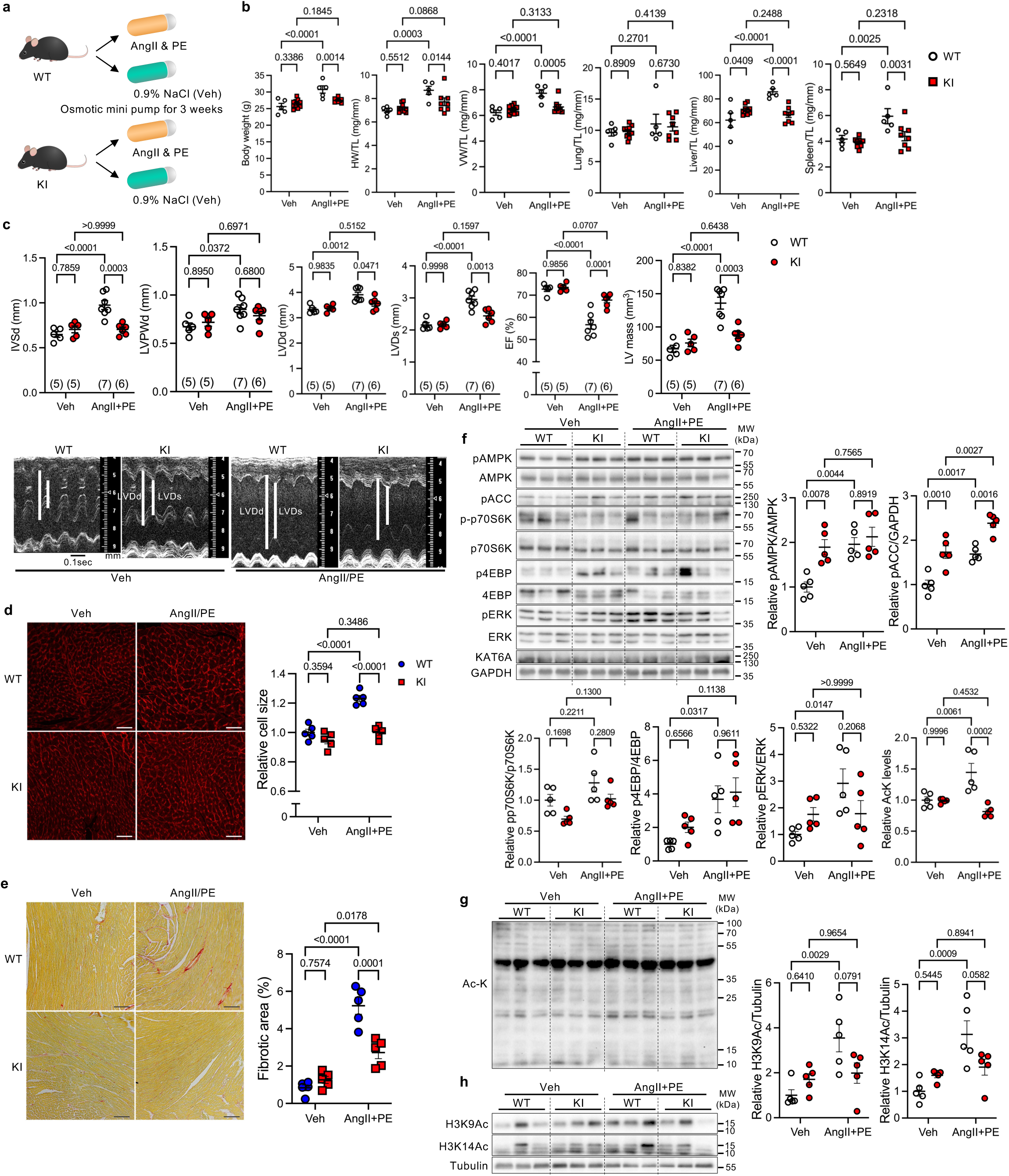
KAT6A-K816R acetylation-resistant knock-in mice exhibit resistance to hypertrophic stimuli in male mice. **a**, Schematic representation of the experimental model. Male knock-in (KI) and WT mice at 10 weeks of age were subjected to chronic angiotensin II (AngII) and phenylephrine (PE) treatment or vehicle control (Veh, 0.9% sodium chloride) via osmotic mini pumps for 3 weeks. **b**, Body weight and organ weights of KI and WT mice after 3 weeks of treatment and normalized to tibia length (TL). Measured weights include heart weight (HW), ventricular weight (VW), lung weight, liver weight, and spleen weight. WT (n=5), KI (n=10 for Veh, n=8 for AngII+PE). **c**, Echocardiographic parameters, including septal wall thickness at end-diastole (IVSd), LV posterior wall thickness at end-diastole (LVPWd), LV end-diastolic diameter (LVDd), LV end-systolic diameter (LVDs), ejection fraction (EF), and LV mass in KI and WT mice after 3 weeks of treatment. Representative M-mode echocardiography images are shown in the lower panel. **d**, Wheat germ agglutinin (WGA) staining of heart sections from KI and WT mice after 3 weeks of treatment. Quantitative analysis of relative cardiomyocyte size (n=5). Scale bar, 50 μm. **e**, Picric Acid Sirius Red (PASR) staining of heart sections from KI and WT mice after 3 weeks of treatment. Quantitative analysis of fibrotic areas (n=5). Scale bar, 100 μm. **f**-**h**. Immunoblots showing AMPK and mTOR signaling pathways (f), overall lysine acetylation (K-Ac) (g), and histone H3K9 and H3K14 acetylation (h) in the hearts of homozygous KI and WT mice after 3 weeks of treatment. Right panels show densitometric analyses of relative protein expression (n=5). Two-way ANOVA followed by Tukey’s multiple comparison test. *N* represents biologically independent replicates. *p* values are shown in the figure. Data are mean ± SEM. Source data are provided as a Source Data file.

RNA-sequencing, proteomics, and physiological analyses indicate the crucial role of AMPK signaling as downstream of KAT6A-K816 acetylation in regulating cardiomyocyte size and energy metabolism. Hypertrophic stimuli at least initially enhance AMPK activity due in part to increases in the ratio of AMP/ATP and the expression of AMPK α1 subunit^19^, which serves as cardioprotection against hemodynamic stress. Consistent with the previous finding^19^, phosphorylation of AMPK and ACC was increased in WT mouse hearts in response to hypertrophic stimuli (Fig 6f). AMPK activity was high at baseline and was further activated in KAT6A-K816R knock-in mouse hearts after hypertrophic stimuli (Fig 6f). Although cardiac hypertrophy was attenuated in the knock-in mice, the levels of phosphorylated p70S6K and 4EBP were similar between two groups after hypertrophic stimuli (Fig. 6f). In contrast to mTOR pathway, stress-induced ERK activation was suppressed in KAT6A-K816R knock-in mice (Fig. 6f). Regarding lysine acetylation, similar to fasting conditions, angiotensin II and phenylephrine treatment increased the levels of acetylation in total lysine residues, H3K9, and H3K14 in the hearts of WT mice (Fig. 6g,h). However, notably, these changes were not observed in the hearts of KAT6A-K816R knock-in mice (Fig. 6g,h).

### Activation of AMPK signaling by neurohumoral stimuli was suppressed in adult cardiomyocytes isolated from KAT6A-K816R knock-in mice

To understand the underlying mechanism of the discrepancies in the effects of KAT6A-K816 acetylation on cell growth and AMPK signaling between *in vivo* mouse hearts and *in vitro* cardiomyocytes, we considered several contributing factors: 1) overexpression versus loss-of-function models; 2) metabolic differences between neonatal and adult cardiomyocytes; and 3) cell-type-specific roles. To address these possibilities, we isolated adult cardiomyocytes from KAT6A-K816R knock-in and WT mice and treated them with angiotensin II and phenylephrine for 30 minutes to evaluate the cell type-specific effects of KAT6A-K816 acetylation on AMPK signaling (Fig. 7a). Cardiomyocytes isolated from WT mice exhibited greater AMPK phosphorylation in response to angiotensin II and phenylephrine treatment compared to those isolated from KAT6A-K816R knock-in mice (Fig. 7b). In line with this, phosphorylation of p70S6K was more pronounced in KAT6A-K816R cardiomyocytes than in WT cardiomyocytes following stimulation (Fig. 7b). H3K9 acetylation decreased in WT cardiomyocytes, while H3K14 acetylation decreased in KAT6A-K816R knock-in cardiomyocytes, in response to hypertrophic stimuli (Fig. 7c,d). Baseline AMPK activity was comparatively lower in KAT6A-K816R knock-in adult cardiomyocytes *in vitro* than in KAT6A-K816R knock-in mouse hearts *in vivo*. Both were relatively lower than baseline AMPK activity observed in neonatal cardiomyocytes, suggesting that K816 acetylation differentially regulates AMPK activity across different cell types and stages of maturation. Nonetheless, hypertrophic stimuli-induced AMPK activation was similarly suppressed in both KAT6A-K816R mouse hearts and KAT6A-K816R adult cardiomyocytes. This underscores the metabolic distinctions between neonatal and adult cardiomyocytes and the potential influence of experimental models (gain- or loss-of function). These findings highlight the intricate regulation of AMPK activity via KAT6A-K816 acetylation in a cell-type-specific manner and reflect metabolic disparities across cardiomyocyte maturation stages. Collectively, these *in vivo* and *in vitro* experiments emphasize the critical role of KAT6A in regulating AMPK activity through K816 acetylation and its pivotal function in cardiomyocyte stress responses.

**Fig. 7:**
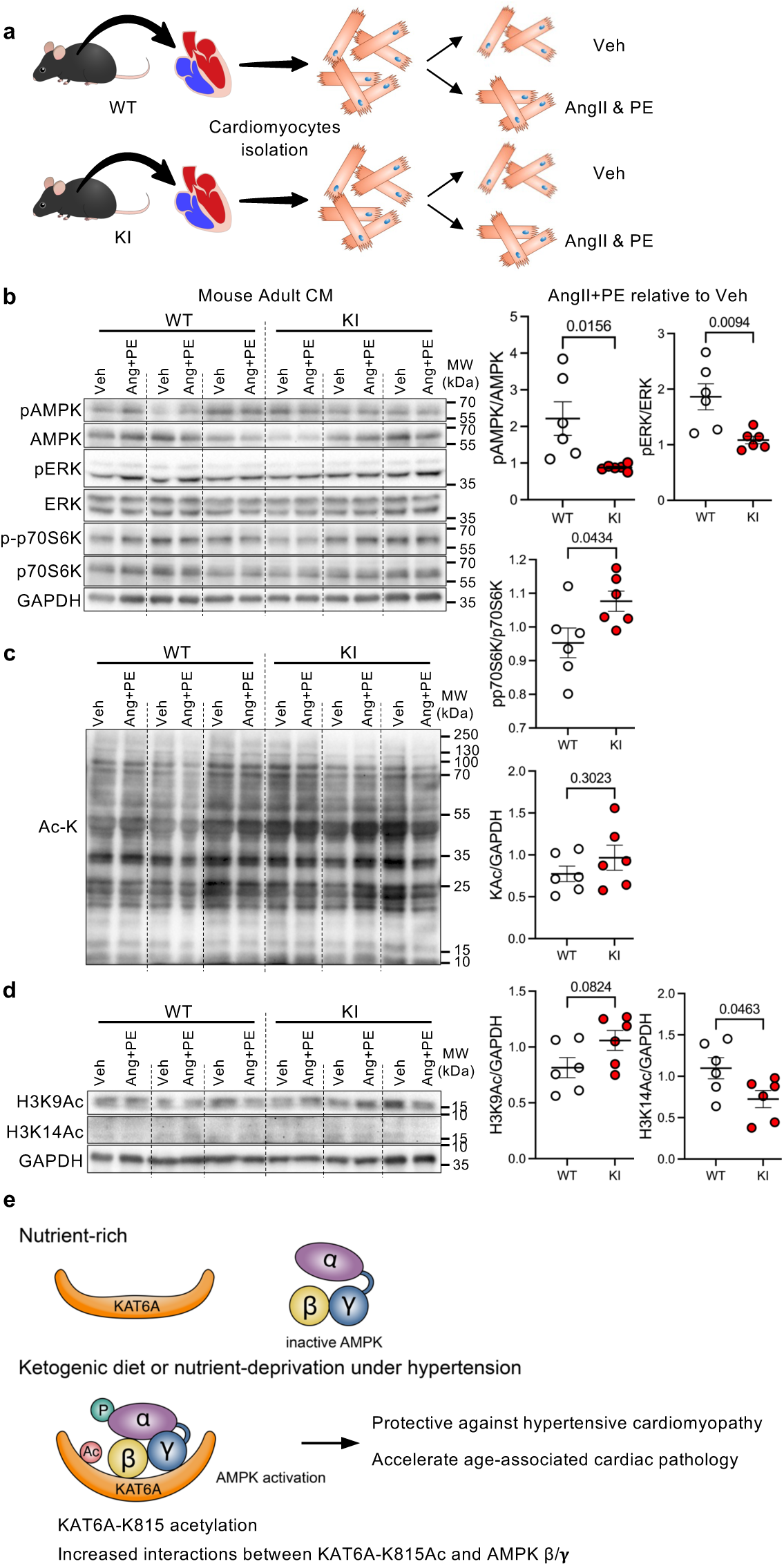
Inhibition of KAT6A acetylation suppresses hypertrophic stimuli-induced AMPK activation in adult mouse cardiomyocytes. **a**, Schematic representation of the experimental approach Adult cardiomyocytes were isolated from KAT6A-K816R knock-in and age-matched WT mice at 12 weeks of age. Cardiomyocytes from the same mice were divided into two groups and treated with angiotensin II (AngII) and phenylephrine (PE) or PBS (Veh) for 30 minutes. **b-d**, Immunoblots showing the AMPK, ERK, and mTOR signaling pathways (b), overall lysine acetylation (K-Ac) (c), and histone H3K9 and H3K14 acetylation (d) in adult cardiomyocytes isolated from 12-week-old homozygous KI and WT mouse hearts in response to AngII and PE treatment for 30 minutes. Right panels show densitometric analysis of relative protein expression (n=6). **e**, Schematic representation of proposed mechanism. Acetylation of KAT6A at K816 enhances its interaction with AMPK regulatory subunits β/γ, promoting AMPK activation in neonatal cardiomyocytes, thereby protecting against hypertensive stress. Unpaired *t* test. *N* represents biologically independent replicates. *p* values are shown in the figure. Data are mean ± SEM. Source data are provided as a Source Data file.

### Lack of KAT6A-K816 acetylation accelerates aging-associated cardiac decline

Acetyltransferases play critical roles in regulating senescence^16^. To investigate the long-term effects of chronic AMPK activation resulting from a lack of KAT6A-K816 acetylation, we monitored KAT6A-K816R knock-in mice up to 28 weeks of age. In male knock-in mice, body weight was slightly but significantly increased compared to WT mice, whereas no significant differences were observed in females (Extended Data Fig. 2b). Similarly, hypertrophy indices -including heart weight, ventricular weight, and LV weight-were elevated in older male, but not female, knock-in mice (Extended Data Fig. 2b). Moreover, echocardiography analysis revealed mild yet significant systolic dysfunction and cardiac remodeling in aged male KAT6A-K816R knock-in mice, effects that were not observed in females (Extended Data Fig. 2c). Furthermore, activation of AMPK signaling and increased acetylation levels of overall lysine residues, as well as histone H3K9 and H3K14, were more pronounced in aged male knock-in mice (Extended Data Fig. 2d,e). While histone H3K9 and H3K14 acetylation and overall lysine acetylation were also elevated in aged knock-in female mice, AMPK signaling was not activated (Extended Data Fig. 2f,g). These findings suggest that the chronic activation of AMPK signaling through the inhibition of KAT6A-K816 acetylation accelerates cardiac aging in male mice, highlighting a sex-specific impact of KAT6A-K816 acetylation on AMPK signaling and age-associated cardiac decline.

## DISCUSSION

In this study, we demonstrate that ketone bodies, such as through exogenous β-hydroxybutyrate administration or dietary intake of a KD, enhance KAT6A acetylation at K815 (corresponding to K816 in mice), which governs heart size and cardiac function in a sex-dependent manner. Our findings reveal that KAT6A-K815 acetylation facilitates its interaction with the regulatory β and γ subunits of AMPK, thereby fine-tuning AMPK signaling. This regulation influences cardiomyocyte size and mitochondrial energetics, providing a cardioprotective mechanism against hypertensive cardiomyopathy in a manner contingent upon cell type and maturation stage of cardiomyocytes (Fig. 7e). These findings establish KAT6A as a pivotal regulatory hub that connects nutrient status to the control of cell size and energy homeostasis, orchestrated through K815 acetylation and the mediation of AMPK signaling pathways.

KATs represent a highly diverse group of enzymes that catalyze the transfer of acetyl groups from acetyl-CoA to lysine residues. KAT6A, also known as monocytic leukemia zinc finger protein (MOZ), is a member of the MYST family of KATs, exhibiting a preference for acetylating histone H3K9 and H3K14, as well as various non-histone proteins^16^. This acetylation activity promotes cell cycle progression and the proliferation of hematopoietic and neural stem cells. KAT6A-deficient mice exhibit embryonic lethality around day 15, with a severe deficit in hematopoietic stem cells^20^. Mutations in the *KAT6A* gene result in the KAT6A syndrome, characterized by intellectual disability, microcephaly, and cardiac defects^17^. Additionally, pharmacological inhibition of KAT6A and KAT6B using small-molecule compounds WM-1119, WM-8014, or PF-07248114 induces cellular senescence in cancer cells, demonstrating anti-tumor effects in mouse models^18^ and holding potential for human applications^21^. Although KAT6A undergoes various post-translational modifications, their functional roles and significance, particularly K815 acetylation, have remained unexplored. In this study, we reveal that KAT6A-K815 acetylation functions as a novel regulatory mechanism that operates independently of its acetyltransferase activity, regulating cell growth and mitochondrial function in cardiomyocytes. It would be valuable to further investigate the roles of KAT6A acetylation in other cell types, such as neural and cancer cells. Notably, previous studies have reported significantly increased KAT6A expression in endothelial and smooth muscle cells in human abdominal aortic aneurysms^22^, reinforcing the cell type-specific role of KAT6A-K815 acetylation observed in the hearts. These findings open avenues for exploring the broader biological implications of KAT6A across various tissues and pathological contexts.

AMPK is a central regulator of energy metabolism and mitochondrial homeostasis by sensing the ATP to AMP ratio to maintain a balance between catabolic and anabolic processes and also regulating autophagy through phosphorylation of key autophagy machinery proteins^23^. AMPK operates as heterotrimeric complexes composed of catalytic α-subunits and regulatory β- and γ-subunits. The γ-subunits bind nucleotides (ATP, ADP, or AMP), inducing conformational changes in the regulatory β- and γ-subunits and modulating AMPK activity in conjunction with phosphorylation at threonine 172 (Thr172). Additionally, AMPK senses the availability of glucose, glycogen, and long-chain fatty acyl-CoA esters, thereby exerting broad control over carbohydrate and lipid metabolism^24, 25^. Our study reveals that ketone bodies promote KAT6A K815 acetylation, which in turn enhances AMPK signaling in cardiomyocytes. These findings suggest a novel regulatory mechanism of AMPK activity mediated by ketone-induced KAT6A acetylation, offering new insights into the interplay between nutrient sensing, acetylation, and metabolic regulation in the heart.

AMPK exhibits diverse subcellular localizations, with its catalytic α-subunit containing nuclear export and import sequences that enable shuttling between the cytoplasm and nucleus, thereby regulating its kinase activity in specific subcellular compartments^26^. Notably, the AMPK γ2 subunit has been shown to translocate into the nucleus during glucose deprivation in COS7 cells^27^. Given the predominant nuclear localization of KAT6A and the minimal effect of K815 acetylation on its distribution, it is likely that acetylated KAT6A at K815 interacts with AMPK β/γ regulatory subunits in the nucleus, which influences the configuration of the AMPK heterotrimeric complex, promoting AMPK phosphorylation and stabilizing phosphorylated Thr172 by shielding it from dephosphorylation. It is plausible that the differential subcellular localization and expression levels of AMPK α/β/γ subunits across cell types, species, and stress conditions contribute to the cell type-specific role of KAT6A-K815 acetylation. Importantly, while KAT6A is known to acetylate non-histone proteins such as p53, our proteomic analysis did not detect AMPK acetylation under conditions where KAT6A is acetylated in the hearts of mice fed a KD. Thus, it is unlikely that KAT6A directly acetylates AMPK to enhance its activity. Instead, our findings suggest that KAT6A functions as a scaffolding protein, facilitating AMPK signaling by promoting the active conformation of the AMPK complex.

Ketone bodies have multiple roles: 1) generating acetyl-CoA through ketolysis to fuel ATP production in mitochondria; 2) functioning as signaling molecules through chromatin modification and non-histone protein acetylation; and 3) acting as endogenous inhibitors of lysine deacetylases (KDACs)^28, 29, 30, 31^. Indeed, ketone bodies increase lysine acetylation levels in both whole lysates and subcellular fractions, including mitochondria, cytosol, and nuclei, in the heart *in vivo* and in cardiomyocytes *in vitro*^8^. Additionally, local changes in acetyl-CoA levels, such as nuclear accumulation, can modulate specific cellular functions^32^. For example, nuclei at the tissue surface exhibit elevated levels of acetyl-CoA synthase, which generates acetyl-CoA to drive histone acetylation in the epithelial tissue of the *Drosophila* wing disc^33^. Our current findings elucidate the role of ketone body in regulating AMPK activity through KAT6A-K815 acetylation. We have previously shown that ketone bodies suppress mTOR signaling in cardiomyocytes^8^; however, modulation of KAT6A-K815 acetylation has minimal impact on the phosphorylation levels of downstream targets such as 4EBP and p70S6K. This discrepancy suggests that ketone body-mediated suppression of mTOR signaling may occur independently of KAT6A acetylation, underscoring the complexity of ketone body actions in cellular signaling and metabolic regulation.

In addition to the response of cardiomyocytes to ketone bodies, metabolic maturation and remodeling unique to the heart have been extensively studied. During postnatal development, cardiac metabolism undergoes a significant shift from glycolysis to fatty acid oxidation, driven by adaptions to environmental changes in nutrient availability, oxygen levels, gut microbiota population, and hemodynamics^34, 35^. Similarly, the expression pattern of AMPK isoforms varies across developmental stages and disease states^36^, aligning with our findings that adult cardiomyocytes isolated from mouse hearts exhibit distinct AMPK activity compared to neonatal cardiomyocytes *in vitro* and mouse hearts *in vivo* in WT. These metabolic characteristics of the heart may underpin the maturation stage-dependent mechanism of KAT6A-K815 acetylation. Furthermore, adenovirus-mediated KAT6A expression increases histone H3 acetylation independently of K815 mutant types, emphasizing the effect of KAT6A expression levels on histone acetylation. Conversely, homozygous KAT6A acetylation-resistant knock-in mice almost completely abolish KAT6A-K815 acetylation. This disruption may either fall outside the physiological range of acetylation levels or represent a compensatory negative feedback mechanism that enhances AMPK activity *in vivo*. These observations highlight the nuanced roles of KAT6A-K815 acetylation in the regulation of cardiac metabolism and function.

In conclusion, our study demonstrates that ketone bodies enhance KAT6A acetylation at K815, facilitating its interaction with AMPK regulatory subunits and thereby stimulating AMPK signaling in cardiomyocytes. These findings highlight the pivotal role of KAT6A acetylation as a central mediator linking ketone availability to metabolic adaptation, regulating cardiomyocyte growth and mitochondrial energy metabolism through AMPK in a manner dependent on sex, cell type, and maturation stage.

## METHODS

### Mice

KAT6A K816R knock-in mice (C57BL/6J background) were generated using CRISPR-Cas9. C57BL/6J embryos were microinjected with a mixture containing High Fidelity Cas9 protein (IDT), an sgRNA (MilliporeSigma) and ssODN (IDT) which contained homology arms and the K816R mutation. The sgRNA used was TTGTTTTCCTTAATTAGGTG<u>GGG</u> (PAM is underlined). The K816R donor oligo sequence (non-coding strand) was CTTCTTGTCAGTTTCTATTAAAGAGGAAGACTCCTGATCTTTTTTCTCCCGAGACACAGAC**c**T **T**CCCACCTAATTAAGGAAAACAAAACAAACAAAAACAACATTGATTTTTTTTCAGTCTGCTT GGTTAAG (K816R sequence change in lower case bold, PAM change in upper case bold). Founders were screened by PCR and digested with restriction enzyme HpyAV (NEB) then confirmed by NGS sequencing (Azenta Life Sciences). Confirmed founders were bred and progeny were screened using PCR primers: KAT6A-A GTGGCCAAAGGGAAAGAGTATTCAG and KAT6A-B GGATAGTGTCTCATCCACAAGCATC and presence of restriction site or by Sanger sequencing of PCR products. Genotyping was performed by Sanger Sequencing of PCR primers using a forward primer (5’-CAAAGGGAAAGAGTATTCAGGGT -3’) and a reverse primer (5’ - ACTTTTCCTCGCAGTCTCCA -3’ for sequencing) to identify the expected mutation (K > R) as shown in Extended Data Figure 2. The mutant offspring were backcrossed into the C57BL/6J background for more than 6 generations. Male C57BL/6J wild-type mice were purchased from Jackson Labs (Strain # 000664) at 5-8 weeks of age to backcross the knock-in mice. Mice of both sexes were used except experiments of hypertrophic stimulation. Mice were housed in a temperature and humidity-controlled environment within a range of 21°C - 23°C with 12-hour light/dark cycles, in which they received food and water *ad libitum*. We used age-matched male mice in all animal experiments. The sample size required was estimated to be n = 5-8 per group according to the Power analysis based upon previous studies examining the effects of pressure overload on cardiac hypertrophy and hypertrophic signaling. All protocols concerning the use of animals were approved by the Institutional Animal Care and Use Committee at Rutgers New Jersey Medical School and all procedures conformed to NIH guidelines (Guide for the Care and Use of Laboratory Animals). Handling of mice and euthanasia with CO_2_ in an appropriate chamber were conducted in accordance with guidelines on euthanasia of the American Veterinary Medical Association. Rutgers is accredited by AAALAC International, in compliance with Animal Welfare Act regulations and Public Health Service (PHS) Policy on Humane Care and Use of Laboratory Animals, and has a PHS Approved Animal Welfare Assurance with the NIH Office of Laboratory Animal Welfare (D16-00098 (A3158-01)).

### Ketogenic diet

Mice were fed a custom diet (Ketogenic diet (D22021702, 7 kcal% Protein and 90% kcal% Fat) or Control diet (D22021701, 10 kcal% Protein and 15 kcal% Fat) (Extended Data Table 8), purchased from Research Diets) ad libitum. Daily *ad libitum* food intake was measured after an initial 2-day acclimation period on a special diet followed by a 5-day experimental period by weighing food provided and remaining every 24 hours and taking an average. The body weight was measured in the late afternoon every week during the special food feeding period. Fasting blood sugar levels were measured after overnight fasting using Aimstrip plus blood glucose meter kit (VWR, #10025-286). Serum β-hydroxybutyrate levels were measured using a β-hydroxybutyrate colorimetric assay kit (Caymen #700190) following the manufacturer’s instructions.

### Hypertrophy induction with Angiotensin II/Phenylephrine treatment

Chronic treatment of hypertrophic agonists (400 μg/kg/day of angiotensin II and 100 mg/kg/day of phenylephrine) or vehicle control (0.9% sodium chloride) was conducted with Alzet osmotic mini-pumps (Alzet, Durect Corporation, Lafeyette Instrument Company, Northern California) for 3 weeks, which were subcutaneously implanted in 9∼10-week-old mice under anesthesia of 2.5% Avertin (10 μl/g body weight) and inhaled 2% isoflurane.

### Echocardiography

Mice were anesthetized using 10 μl/g body weight of 2.5% avertin (Sigma-Aldrich), and echocardiography was performed using ultrasound (Vivid 7, GE Healthcare). It took around 10-20 minutes from the establishment of anesthesia to the completion of echocardiography and 1-2 hours to fully recover from anesthesia after echocardiography. A 13-MHz linear ultrasound transducer was used. Mice were subjected to 2-dimension guided M-mode measurements of LV internal diameter at the papillary muscle level from the short-axis view to measure systolic function and wall thickness, which were taken from at least three beats and averaged. LV ejection fraction was calculated as follows: Ejection fraction = [(LVEDD)^3^ – (LVESD)^3^]/(LVEDD)^3^ x 100. LV mass was calculated as follows: LV mass = 1.05 x [(IVSd+LVDd+LVPWd)^3^ – (LVDd)^3^].

### Cell line

HEK293 cells were purchased from the American Type Culture Collection (CRL-1573) and maintained at 37°C with 5% CO_2_ in Dulbecco’s modified Eagle’s Medium with 10% fetal bovine serum and penicillin/streptomycin. H9C2 cells were purchased from the ATCC and were maintained at 37 °C with 5% CO_2_ in Dulbecco’s modified Eagle’s medium/Nutrient Mixture F-12 supplemented with 10% fetal bovine serum and penicillin/streptomycin.

### Primary rat neonatal cardiomyocytes

Primary cultures of ventricular cardiomyocytes were prepared from 1-day-old Crl:(WI)BR-Wistar rats (Envigo, Somerville) and maintained in culture as described previously^14, 15^. The neonatal rats of both sexes were deeply anesthetized with isoflurane. The chest was opened, and the heart was harvested. A cardiomyocyte-rich fraction was obtained by centrifugation through a discontinuous Percoll gradient. Cardiomyocytes were cultured in complete medium containing Dulbecco’s modified Eagle’s medium/F-12 supplemented with 5% horse serum, 4 μg/ml transferrin, 0.7 ng/ml sodium selenite, 2 g/l bovine serum albumin (fraction V), 3 mM pyruvate, 15 mM Hepes pH 7.1, 100 μM ascorbate, 100 mg/l ampicillin, 5 mg/l linoleic acid, and 100 μM 5-bromo-2’-deoxyuridine (Sigma). Culture dishes were coated with 0.3% gelatin or 2% gelatin for immunofluorescence staining on chamber slides. The following ketone bodies were used; β-hydroxybutyrate (Sigma-Aldrich #H6501) and acetoacetate (SigmaAldrich # A8509). WM-1119 (Tocris Bioscience, #6692) was dissolved in DMSO and used for the indicated concentration.

### Isolation of mouse adult cardiomyocytes

Adult mouse cardiomyocytes were isolated as described previously with a modification^15, 37^. Briefly, the heart of a male mouse was perfused with 12 ml EDTA buffer [130 mM NaCl, 5 mM KCl, 0.5 mM NaH_2_PO_4_, 10 mM HEPES, 10 mM Glucose, 10 mM BDM, 10 mM Taurine, 5 mM EDTA] to stop the beating of the heart. Digestion was achieved using 30 ml perfusion buffer [130 mM NaCl, 5 mM KCl, 0.5 mM NaH_2_PO_4_, 10 mM HEPES, 10 mM Glucose, 10 mM BDM, 10 mM Taurine, 1 mM MgCl_2_] containing 1 mg/ml Collagenase type II (Worthington, LS004177) and 0.05 mg/ml Protease XIV (Sigma-Aldrich, P5147). Cellular dissociation was stopped by addition of 5 ml Perfusion buffer containing 5% FBS and 100 mM BSA-conjugated fatty acid cocktail (palmitic acid: oleic acid: linoleic acid = 2:1:1 and BSA: fatty acid = 1:5). Cell suspension was passed through a 100-μm filter and cardiomyocytes, and non-cardiomyocytes were separated by 4 sequential rounds of gravity settling with calcium reintroduction medium containing 100 mM BSA-conjugated fatty acid cocktail. The non-myocyte fraction was plated on 6-well tissue-culture plate in a humidified tissue culture incubator (37°C, 5% CO_2_). Media was changed to Dulbecco’s modified Eagle’s Medium (Cytiva, SH30243.01) with 10% fetal bovine serum after 24 hours.

### Adenovirus constructs

Recombinant adenovirus vector for overexpression was constructed, propagated and titered as previously described^15, 38, 39^. Briefly, pBHGloxΔE1,3Cre (Microbix), including the ΔE adenoviral genome, was co-transfected with the pDC shuttle vector containing the gene of interest into HEK293 cells. Replication-defective human adenovirus type 5 (devoid of E1) harboring human full length wild type or mutant KAT6A cDNA (Ad-FLAG- or YFP-tagged KAT6A or Ad-KAT6A without tag) was generated by homologous recombination in HEK293 cells. Generation of adenovirus harboring a dominant negative form of the α2 subunit of AMPK has been described^40^. Adenovirus harboring beta-galactosidase (Ad-LacZ) was used as a control.

### Immunoblotting

Cardiomyocyte lysates and heart homogenates were prepared in RIPA buffer containing protease and phosphatase inhibitors (Sigma-Aldrich P8340 and P0044)^8, 14, 15, 38^. Lysates were centrifuged at 16,100 x g at 4°C for 15 minutes. Protein concentrations were determined using a standard BCA assay (Thermo Fisher Scientific, 23227). Total protein lysates (25-45 μg) were incubated with SDS sample buffer (final concentration: 100 mM Tris (pH 6.8), 2% SDS, 5% glycerol, 2.5% 2mercaptoethanol, and 0.05% bromophenol blue) at 95°C for 5 minutes. The denatured protein samples were separated by SDS-PAGE, transferred to polyvinylidene difluoride membranes (Bio-Rad, 1620177) by semi-dry electrophoretic transfer (Bio-Rad), blocked in 2% (w/v) BSA (Fisher, BP9703100) in 1×TBS/0.5% Tween 20 at room temperature for 1 hour, and probed with primary antibodies at 4°C overnight. After washing with 1×TBS/0.5% Tween 20 for 20 minutes, the membranes were incubated with the corresponding secondary antibody at room temperature for 1 hour. After washing with 1×TBS/0.5% Tween 20 for 45 minutes, the membranes were developed with ECL Western blotting substrate (Millipore, WBKLS0500, WBLUC0100), followed by acquisition of digital image with the ChemiDoc MP Imaging System (Bio-Rad). The intensities of Western blot bands were quantified using ImageJ software. Uncropped blotting images with molecular markers are provided in Source Data File.

### Antibodies and reagents

The following commercial antibodies were used at the indicated dilutions: rabbit monoclonal phospho-AMPKα (Thr172) antibody (#2535) (1:3,000), rabbit monoclonal AMPKα antibody (#2603) (1:3,000), rabbit monoclonal phospho-Acetyl-CoA Carboxylase (Ser79) antibody (#3661) (1:3,000), rabbit monoclonal Acetyl-CoA Carboxylase antibody (#3662) (1:2,000), rabbit monoclonal phospho-4E-BP1 (Thr70) antibody (#9455) (1:3,000), rabbit monoclonal 4E-BP1 antibody (#9452) (1:3,000), rabbit monoclonal phospho-p70 S6 Kinase (Thr389) antibody (#9205) (1:3,000), rabbit monoclonal p70 S6 Kinase antibody (#9202) (1:3,000), rabbit monoclonal GAPDH (#5174) (1:5,000), Acetyl-Histone H3 (Lys9) antibody (#9649) (1:3,000), rabbit monoclonal Acetyl-Histone H3 (Lys14) antibody (#7627) (1:2,000), rabbit monoclonal Acetylated-Lysine antibody (#9441), anti-mouse or -rabbit IgG, HRP-linked antibodies (#7076 and #7074) (1:5,000) (Cell Signaling Technology); mouse monoclonal α-tubulin antibody (T6074) (1:5,000), mouse monoclonal FLAG M2 antibody (F1804) (1:4,000) (Sigma-Aldrich); Antibodies were diluted in either 2% (w/v) BSA in 1×TBS/0.5% Tween 20 or Immunobooster solution (Takara), depending on the level of background intensity.

### FLAG Pull-down assay

Cardiomyocytes were transduced with adenovirus harboring FLAG-KAT6A-WT, K815R, or K815Q for 2 days. Control FLAG sample was used as a negative control. The cardiomyocytes were collected with RIPA buffer containing protease and phosphatase inhibitors (Sigma-Aldrich) as described previously ^38^. Lysates were centrifuged at 16,100 x g at 4°C for 15 minutes. After protein concentrations were determined using a standard BCA assay, protein lysates were incubated with anti-FLAG M2 Magnetic Beads (Sigma-Aldrich) overnight at 4°C. After FLAG pull-down, the samples were washed with lysis buffer five times and eluted with lysis buffer containing 100 μg/mL 3x FLAG peptide (Sigma-Aldrich, Cat#F4799). KAT6A interacting proteins were separated by SDS-PAGE and stained with Coomassie Brilliant Blue, followed by Mass Spectrometry.

### Mass Spectrometry Sample Preparation and Analysis for protein-protein interaction

The whole four lanes of SDS-PAGE gel samples (Ctrl, KQ, KR and WT) were excised for in-gel trypsin digestion as described^14^. Briefly, the gel bands of each sample were excised into ∼1 cm^3^ pieces and washed four times with 1 μL each of a solution of 30% acetonitrile (ACN) and 70% of 100 mM NH_4_HCO_3_. Subsequently, 200 μL of 25 mM dithiothreitol (DTT) solution was used to reduce disulfides at 55 °C for 30 minutes, and 200 μL of 50 mM iodoacetamide solution was then added to alkylate thiols at room temperature in the dark for 30 minutes. The resulting gel pieces were dehydrated with 200 μL of ACN to remove both DTT and iodoacetamide. For in-gel trypsin digestion, 100 μL of trypsin solution (5 µg/mL in 50 mM NH_4_HCO_3_) was added into each sample and incubated at 37 °C overnight. Resulting peptides were extracted, desalted with Pierce C_18_ spin columns (Thermo Scientific), based on the manufacturer’s protocol, followed by Speed Vac prior to LC-MS/MS analysis on a Thermo Orbitrap Fusion Lumos MS instrument.

Peptides from each sample were reconstituted in Solvent A (consisting of 2% ACN in 0.1% formic acid, FA). Two microliters of peptides from each sample were subjected to LC-MS/MS analysis using an Orbitrap Fusion Lumos Mass Spectrometer coupled with an UltiMate 3000 UHPLC^+^ system (Thermo Scientific). The separation of peptides occurred on an Acclaim PepMap C_18_ column (75 µm × 50 cm, 2 µm, 100 Å) with a 2-hour binary gradient ranging from 2% to 95% of Solvent B (85% ACN in 0.1% FA), at a flow rate of 300 nL/min. Eluted peptides were introduced into the MS system *via* a Nanospray Flex ion source with a spray voltage of 2 kV and a capillary temperature of 275 °C. MS spectra were acquired in positive ion mode using Xcalibur (1.5). Full MS scans were obtained in an m/z range of 375 to 1500 in profile mode, with an AGC value of approximately 1E6. Subsequent to each full MS scan, the data-dependent MS/MS mode was used to analyze the ions with charge states ranging from 2^+^ to 7^+^. The isolation window of 2 m/z was used for MS/MS analysis and the higher energy collision dissociation (HCD) was used for fragmentation with a normalized collision energy of 30%. The AGC for MS/MS analysis was set to 5E4 and a dynamic exclusion time of 45 seconds was implemented. The MS/MS spectra were searched against a Swissprot rat database using Sequest search engines on Proteome Discoverer (V2.3) platform. The protein false discovery rate is less than 1% as listed in the attached Table. The Ratio of (KQ/Ctrl), (KR/Ctrl) and (WT/Ctrl) were calculated using spectra counting method.

### Mass Spectrometry Sample Preparation and Analysis for acetylation protein identification

Heart tissues from four mice on a ketogenic diet and four on a control diet were collected and pooled for identification of protein acetylation. The samples were homogenized in the lysis buffer containing 8M urea, 100 mM ammonium bicarbonate, and 1× protease inhibitor cocktail, followed by high-speed centrifugation. Protein concentrations were determined using the Branford Assay. Fifteen milligrams of protein from each sample were used for analysis. We performed in-solution trypsin digestion. The resulting peptides were desalted using a C18 column. A small aliquot of the samples was used for protein expression analysis, while the rest of samples was used for the enrichment of acetylation peptides using the PTMScan Acetyl-Lysine Motif [Ac-K] Kit (Cell Signaling Technology). The enriched peptides were analyzed by LC-MS/MS on a Q Exective MS instrument. MS/MS spectra were searched against the Uniprot mouse database using the Sequest search engine, with a false discovery rate for protein and peptide identification of less than 1%, as listed in the attached Tables.

### RNA-Seq Library Preparation, Sequencing, and Data Analysis

Total RNA was isolated from rat neonatal cardiomyocytes using TRIzol (Invitrogen). Isolated RNA was checked for integrity on an Agilent Bioanalyzer 2100; samples with RNA integrity number >7.0 were used for subsequent processing. Total RNA was subjected to two rounds of poly(A) selection using oligo-d(T)25 magnetic beads (New England Biolabs). A paired-end (strand specific) cDNA library was prepared using the NEB Next Ultra-directional RNA-seq protocol. Briefly, poly(A)+ RNA was fragmented by heating at 94°C for 10 minutes, followed by reverse transcription and second strand cDNA synthesis using the reagents provided in the NEB Next kit. End-repaired cDNA was then ligated with double stranded DNA adapters, followed by purification of ligated DNA with AmpureXP beads. cDNA was then amplified by PCR for 15 cycles with a universal forward primer and a reverse primer with bar code. The sequencing of the cDNA libraries was performed on the Illumina NextSeq 500 platform (Illumina, San Diego, CA) using the single-read 1×75 cycles configuration. The raw reads files have been deposited in the NCBI Gene Expression Omnibus.

Raw reads were quality trimmed using Trimmomatic-0.39 with leading and trailing Q score 20, minimum length 35 bp. The cleaned reads were mapped to *Rattus norvegicus* genome Rnor_6.0 using HISAT2 (Version 2.2.1). The reference genome sequence and annotation files were downloaded from ENSEMBL. The aligned read counts were obtained using htseq-count as part of the package HTSeq-0.6.1. The Bioconductor package edgeR (Version 3.18.1 with limma 3.32.10) was used to perform the differential gene expression analysis under R environment, R version 4.1.1. Expression patterns of regulated genes were graphically represented in a heat map. Hierarchical clustering was performed to group genes with similar features in the expression profile. Normalized expression data were also analyzed with GSEA version 4.1.0 software using the JAVA program (Broad Institute, Cambridge, MA). All gene sets were obtained from the Molecular Signatures Database version 7.3 distributed on the GSEA Web site.

### Immunohistochemistry

The heart tissues were washed with PBS, fixed in 4% paraformaldehyde overnight, embedded in paraffin, and sectioned at 10-µm thickness onto a glass slide. After de-paraffinization, sections were stained with wheat germ agglutinin (WGA) for evaluation of the cross-sectional area of cardiomyocytes, or Picric Acid Sirius Red (PASR) for fibrosis. The myocyte cross-sectional area was measured from images captured from wheat germ agglutinin (WGA)-stained sections. The tissues were observed under a fluorescence microscope (BX51, Olympus). The outline of 100–200 myocytes was traced in each section using ImageJ software (NIH).

### Cardiomyocyte cell size measurement

Rat neonatal cardiomyocytes were cultured on coverslips, washed with PBS twice, fixed with 3.7% paraformaldehyde for 15 minutes, and washed with PBS three times. Samples were permeabilized with PBST (0.5% Triton-X in PBS) for 15 min, and blocked in 5% BSA, 5% goat serum in PBST for 30 minutes at 37°C. Cardiomyocytes were stained with Alexa Fluor 555 phalloidin (Thermo Fisher Scientific, A34055). Samples were washed with PBS and mounted on glass slides with mounting medium (VECTASHIELD, Vector Laboratories). Cells were observed under a fluorescence microscope (Eclipse T*i*, Nikon) with Nikon NIS-Elements imaging software. Cell size was measured for 25 - 30 cells for each condition in each experiment and the mean value was taken as representative of the experiment. This experiment was performed independently five times (n = 5).

### Seahorse experiment

The oxygen consumption rate (OCR, pmol/min) in cultured rat neonatal ventricular cardiomyocytes was measured using a Seahorse XFe96 Extracellular Flux Analyzer (Seahorse Bioscience) according to the Installation and Operation Manual from Agilent Technologies. Cardiomyocytes were plated at a density of 30,000 cells/well in 96-well Seahorse assay plates except for background correction wells (Agilent, 102416-100). Cardiomyocytes were transduced with adenovirus for 48 hours prior to measurement. One hour prior to the beginning of measurements, the medium was replaced with XF base medium (Agilent, 103335-100) supplemented with 5.5 mM glucose (Sigma-Aldrich, G8270), 1 mM pyruvate (Agilent, 103578-100) and 2 mM L-glutamine (Agilent, 103579-1001), and cardiomyocytes were incubated for 1 hour in a 37°C incubator without CO_2_. Oxygen consumption rate (OCR) was measured three times at baseline, followed by sequential injection into each well with 2 µM oligomycin (Sigma-Aldrich, O4876) to measure the ATP-linked OCR, 5 µM Carbonylcyanidep-triflouromethoxyphenylhydrazone (FCCP) (Sigma-Aldrich, C2920), an uncoupler, to determine maximal respiratory capacity, and 2 µM rotenone (Sigma-Aldrich, R8875) mixed with 2 µM antimycin A (Sigma-Aldrich, A8674) to determine the non-mitochondrial respiration. For the measurement of fatty acid oxidation rate, etomoxir (100 µM, Sigma-Aldrich E1905), a specific Cpt1 inhibitor, was added to assess the degree of fatty acid oxidation.

### Quantitative RT-PCR

Total RNA was prepared from mouse hearts using TRIzol reagent (Thermo Fisher Scientific, 15596-018) according to the manufacturer’s instructions^14^. Complementary DNA (cDNA) was synthesized by reverse transcription using 480 ng total RNA with PrimeScript RT Master Mix (Takara, RR036). Using Maxima SYBR Green qPCR master mix (ThermoFisher Scientific, K0253), real-time RT-PCR was performed under the following conditions: 95°C for 3 minutes; 40 cycles of 95°C for 10 seconds, 58°C for 15 second, 72°C for 45 seconds; and a final elongation at 72°C for 10 minutes. Relative mRNA expression was determined by the ΔΔ-Ct method normalized to the ribosomal RNA (18S) level. The following oligonucleotide primers were used: NPPA, sense 5’-ATGGGCTCCTTCTCCATCAC-3’ and antisense 5′-ATCTTCGGTACCGGAAGCTG-3′; NPPB, sense 5′-AAGTCCTAGCCAGTCTCCAGA-3′ and antisense 5′-GAGCTGTCTCTGGGCCATTTC-3′; MYH7, sense 5′-GCCAACACCAACCTGTCCAAGTTC-3′ and antisense 5′-TGCAAAGGCTCCAGGTCTGAGGGC-3′; MYH6, sense 5’-GGAAGAGTGAGCGGCGCATCAAGG -3’ and antisense 5’-CTGCTGGAGAGGTTATTCCTCG -3’; 18S rRNA, sense 5′-CGCGGTTCTATTTTGTTGGT-3′ and antisense 5′-AGTCGGCATCGTTTATGGTC-3′.

### Viability and mitochondrial function of cardiomyocytes

Tetramethylrhodamine methyl ester (TMRM) (ThermoFisher Scientific) was used to determine the level of mitochondrial functionality according to the manufacture’s protocol (dilution of the 1000x concentrated stock solution in culture medium). Cells were observed under a fluorescence microscope (Eclipse T*i*, Nikon) with Nikon NIS-Elements imaging software. Intensity of the signals was measured for 8 - 10 cells expressing a YFP-KAT6A mutant (K816R or K816Q) compared to cells without YFP expression (control) for each condition in each experiment and the mean value was taken as representative of the experiment. This experiment was performed independently four times.

## STATISTICS

All values are expressed as mean ± SEM. Statistical analyses were carried out by 2-tailed unpaired Student *t* test for 2 groups or one- or two-way ANOVA followed by the Tukey post-hoc analysis for 3 groups or more unless otherwise stated. If the data distribution failed normality by the Shapiro-Wilk test or Kolmogorov-Smirnov test, the Mann-Whitney *U* test for 2 groups was performed. The statistical analyses used for each figure are indicated in the corresponding figure legends. Survival curves were plotted by the Kaplan-Meier method, with statistical significance analyzed by log-rank test. Statistical analyses were conducted using Prism 10 (GraphPad Software). All experiments are represented by multiple biological replicates or independent experiments. The number of replicates per experiment are indicated in the legends. All experiments were conducted using at least two independent experimental materials or cohorts to reproduce similar results. No sample was excluded from analysis. A *p* value of less than 0.05 was considered significant.

## DATA AVAILABILITY

All data generated or analyzed during this study are included in this published article and its Supplementary Information. RNA-sequencing data have been deposited at GEO and are publicly available as of the date of publication (Accession numbers: GSE286926). The mass spectrometry data have been deposited to the ProteomeXchange Consortium via the PRIDE partner repository. Source data are provided with this paper. Other data and study materials are available from the corresponding author on reasonable request.

## Acknowledgements

We thank Dr. Junichi Sadoshima (Rutgers New Jersey Medical School) for generously providing the adenovirus-harboring dominant-negative AMPK α2. This study was supported in part by U.S. Public Health Service grants HL155766 (M.N.). This work was also supported by American Heart Association Scientist Development grant (17SDG33660358) (M.N.). The mass spectrometry data were obtained from the Rutgers Center for Advanced Proteomics Research, which partially receives funding from NIH grant 1S10OD034300 (H.L.). The KAT6A-K816R knock-in mice were generated by Rutgers Cancer Institute of New Jersey Genome Editing Shared Resource P30CA072720-6852.

## Author contributions

M.N. designed the experiments and wrote the paper; M.N. and M.A.K. conducted the *in vitro* and *in vivo* experiments; M.N. and M.A.K. conducted the animal experiments and analyses; A.I. prepared the histology samples; T.L. and H.L. conducted the mass spectrometry analyses; M.N. conducted the gene expression analyses; P.J.R. generated a KAT6A-K816R knock-in mouse model with a CRISPR-CAS9 system; M.N. generated project resources. All authors reviewed and commented on the manuscript.

## Competing interests

The authors declare no competing interests.

## Extended Data Tables

**Extended Data Table 1**

**Acetylation peptide identified in the hearts of C57BL/6J wild-type mice fed a normal chow (NC) after high blood pressure stress**

Excel file provided separately.

**label>Extended Data Table 2**

**Acetylation peptide identified in the hearts of C57BL/6J wild-type mice fed a ketogenic diet (KD) after high blood pressure stress**

Excel file provided separately.

**Extended Data Table 3**

**Acetylated protein identified uniquely in the hearts of C57BL/6J wild-type mice fed a ketogenic diet (KD) after high blood pressure stress**

Excel file provided separately.

**Extended Data Table 4:** Gene sets enriched in cardiomyocytes transfected with adenovirus-KAT6A compared to adenovirus-LacZ (listed only gene sets with FDR < 25%).

|  | ES | NES | NOM<br>p-val | FDR<br>q-val |
| --- | --- | --- | --- | --- |
| REACTOME_Activated AMPK stimulates fatty acid oxidation in muscle | -0.66 | -1.9 | 0 | 0.067 |
| KEGG_Regulation of autophagy | -0.62 | -1.79 | 0 | 0.173 |
| HOFMANN_Cell lymphoma DN | -0.48 | -1.75 | 0 | 0.196 |

**Extended Data Table 5:**
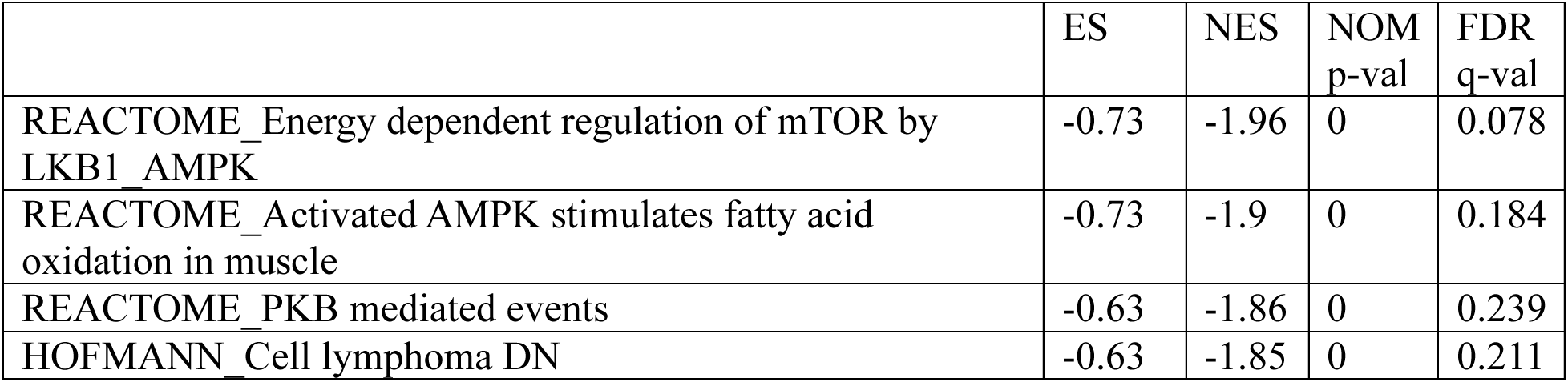
Gene sets enriched in cardiomyocytes transfected with adenovirus-KAT6A-K815Q compared to adenovirus-KAT6A-K815R (listed only gene sets with FDR < 25%).

|  | ES | NES | NOM<br>p-val | FDR<br>q-val |
| --- | --- | --- | --- | --- |
| REACTOME_Energy dependent regulation of mTOR by LKB1 AMPK | -0.73 | -1.96 | 0 | 0.078 |
| REACTOME_Activated AMPK stimulates fatty acid oxidation in muscle | -0.73 | -1.9 | 0 | 0.184 |
| REACTOME_PKB mediated events | -0.63 | -1.86 | 0 | 0.239 |
| HOFMANN_Cell lymphoma DN | -0.63 | -1.85 | 0 | 0.211 |

**Extended Data Table 6**

**Interaction proteins with KAT6A-wild type (WT), K815R, and K815Q mutants, identified by mass spectrometry analysis.**

Control flag sample was used as a negative control.

Excel file provided separately.

**Extended Data Table 7:**
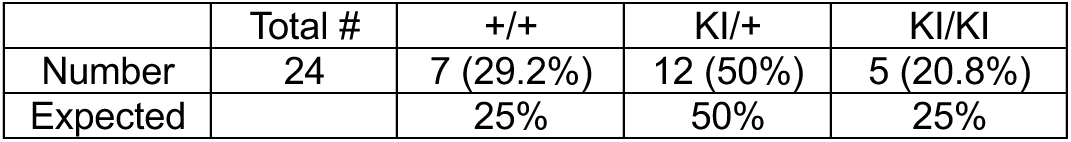
Offspring chart for mice crossed with KAT6A-K816R heterozygous knock-in mice.

**Extended Data Table 8:** The components of custom ketogenic diet and its isocaloric control diet, purchased from Research Diets, Inc.

| Diets (Product #) | D22021701 |  | D22021702 |  |
| --- | --- | --- | --- | --- |
|  | Control |  | Ketogenic |  |
|  | gm% | kcal% | gm% | kcal% |
| Protein | 10 | 10 | 12 | 7 |
| Carbohydrate | 74 | 75 | 5 | 3 |
| Fat | 7 | 15 | 67 | 90 |
| Total |  | 100 |  | 100 |
| Kcal/gm | 4.0 |  | 6.7 |  |
| Ingredient | gm | kcal | gm | kcal |
| Casein, 80 Mesh | 100 | 400 | 70 | 280 |
| L-Cystine | 1.5 | 6 | 1 | 4 |
| Corn Starch | 447.5 | 1790 | 0 | 0 |
| Maltodextrin 10 | 150 | 600 | 21.5 | 86 |
| Sucrose | 150 | 600 | 0 | 0 |
| Cellulose, BM200 | 50 | 0 | 50 | 0 |
| Soybean Oil | 25 | 225 | 25 | 225 |
| Lard | 0 | 0 | 0 | 0 |
| Corn Oil | 0 | 0 | 50 | 450 |
| Cocoa Butter | 0 | 0 | 50 | 450 |
| Hydrogenated Vegetable Oil | 44 | 396 | 280.2 | 2522 |
| Mineral Mix, S10026 | 10 | 0 | 10 | 0 |
| DiCalcium Phosphate | 13 | 0 | 13 | 0 |
| Calcium Carbonate | 5.5 | 0 | 5.5 | 0 |
| Potassium Citrate, 1 H<sub>2</sub>O | 16.5 | 0 | 16.5 | 0 |
| Vitamin Mix, V10001 | 10 | 40 | 10 | 40 |
| Choline Bitartrate | 2 | 0 | 2 | 0 |
| FD&C Yellow Dye #5 | 0.025 | 0 | 0.05 | 0 |
| FD&C Red Dye #40 | 0 | 0 | 0 | 0 |
| FD&C Blue Dye #1 | 0.025 | 0 | 0 | 0 |
| Total | 1025.05 | 4057 | 604.75 | 4057 |

## Extended Data Figures

**Extended Data Fig.1:**
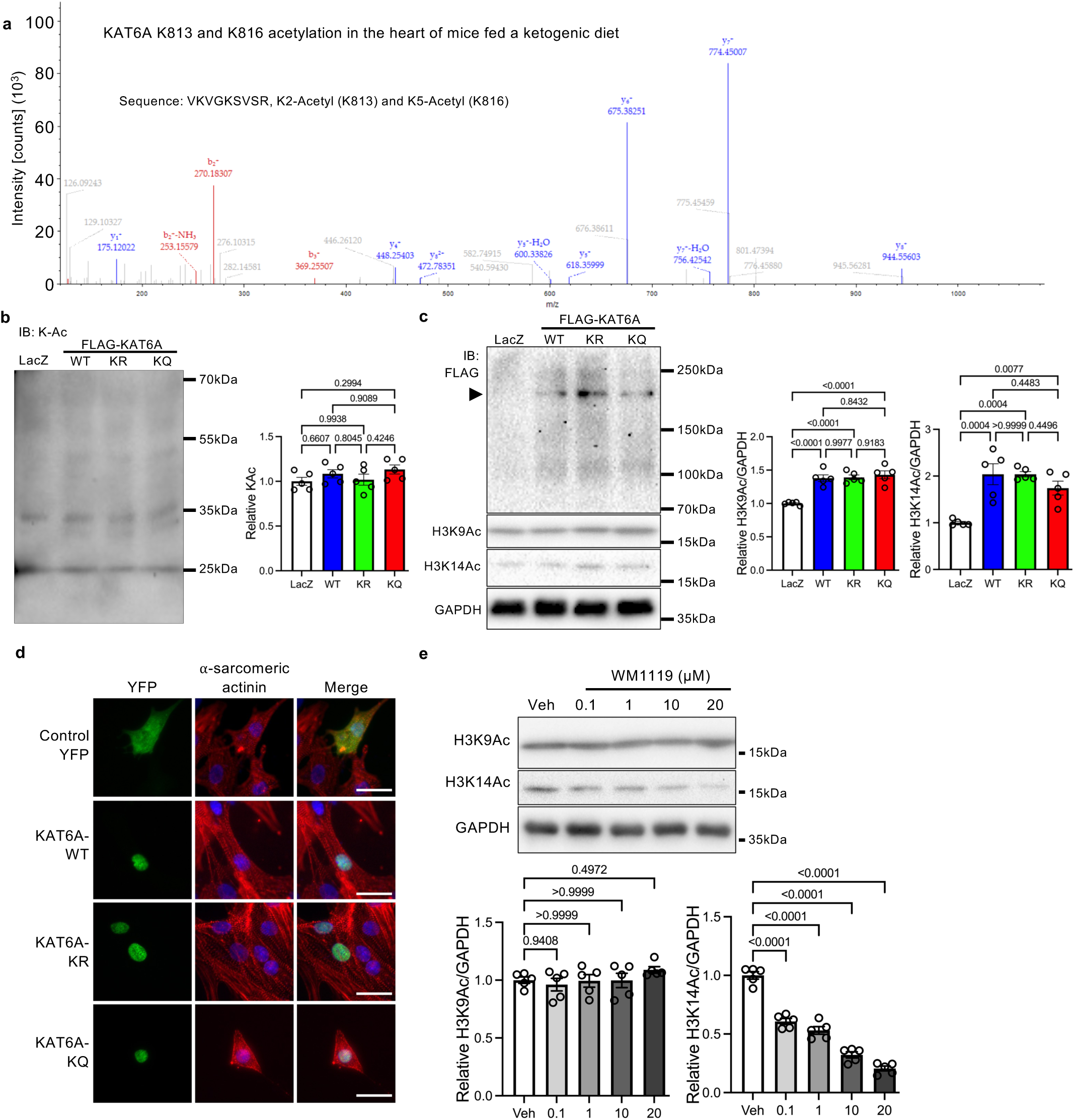
KAT6A is localized in the nucleus and enhances histone H3K9 and H3K14 acetylation independently of K815 acetylation. **a**, Mass spectrometry analysis of KAT6A peptide fragment reveals acetylation at K813 and K816 in the hearts of mice fed a ketogenic diet under high blood pressure. **b-c**, Immunoblots showing overall lysine acetylation (K-Ac) (b) and histone H3K9 and H3K14 acetylation (Ac) (c) in cardiomyocytes expressing FLAG-KAT6A-wild type (WT), K815R (KR), or K815Q (KQ) for 48 hours. The right panels show quantitative analyses of protein expression (n=5). **d**, Immunofluorescent images showing the localization of YFP-KAT6A-WT, -K815R, and - K815Q mutants, co-stained with α-sarcomeric actinin and DAPI. Scale bar, 25 μm. **e**, Immunoblots showing expression levels of histone H3K9Ac and H3K14Ac. Lower panels show quantitative analyses of the expression (n=5). One-way ANOVA followed by Tukey’s multiple comparison test. *N* represents biologically independent replicates. *p* values are shown in the figure. Data are mean ± SEM. Source data are provided as a Source Data file.

**Extended Data Fig.2:**
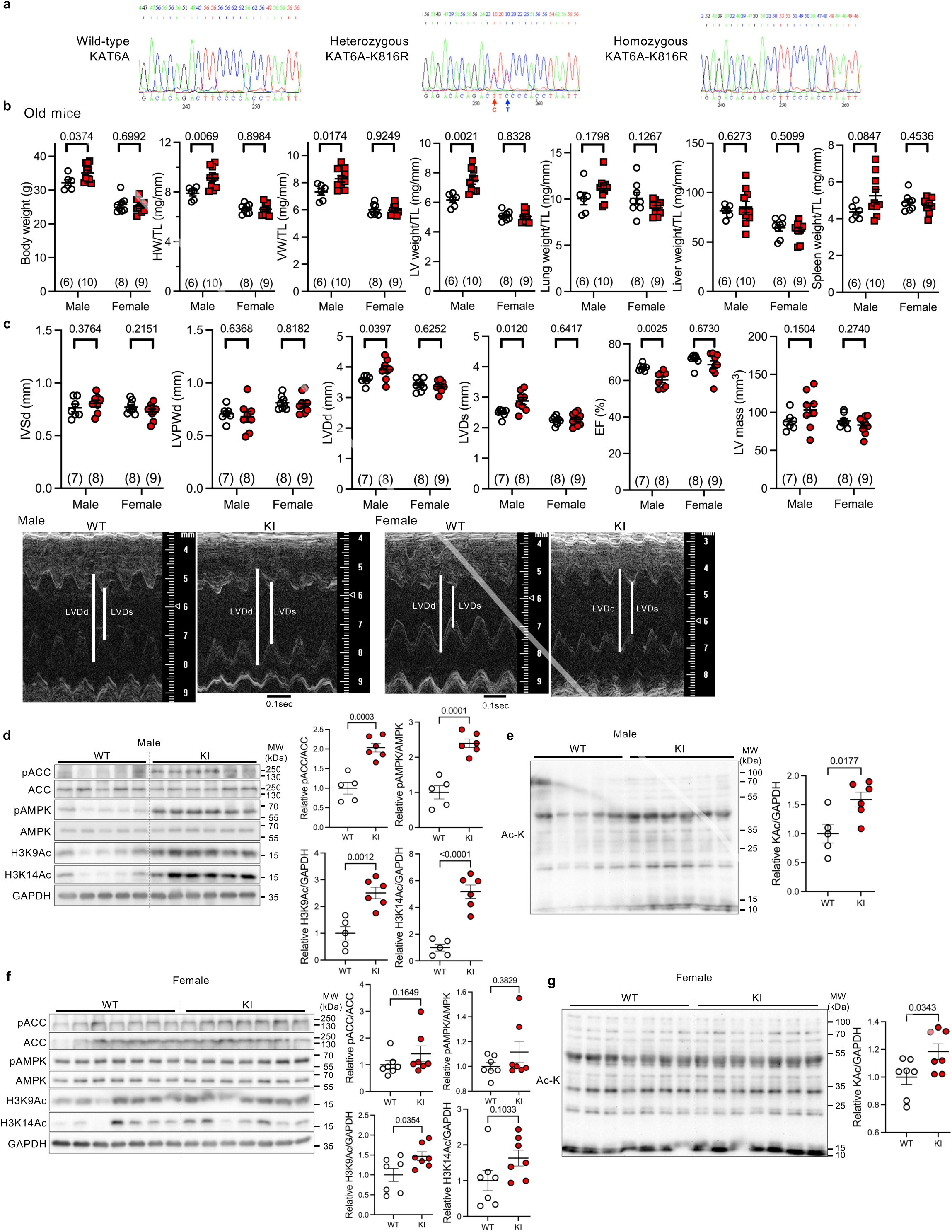
Generation of KAT6A K816R knock-in mice and characterization of their basal phenotypes in advanced age. **a**, Sequencing analysis of the KAT6A region of interest in KAT6A-wild type (WT) and K816R heterozygous, and homozygous knock-in mice. **b**, Body and organ weights of KI and littermate WT mice of both sexes at 24-28 weeks of age, normalized to tibia length (TL). Measured weights include heart weight (HW), ventricular weight (VW), left ventricular (LV) weight, lung weight, liver weight, and spleen weight. Unpaired *t* test within each sex. **c**, Echocardiographic parameters in KI and littermate WT mice of both sexes at 24-28 weeks of age. Measurements include interventricular septal wall thickness at end-diastole (IVSd), LV posterior wall thickness at end-diastole (LVPWd), LV end-diastolic diameter (LVDd), LV end-systolic diameter (LVDs), ejection fraction (EF), and LV mass. Unpaired *t* test within each sex. Representative M-mode echocardiography images are shown in the lower panel. **d**-**g**, Immunoblots showing AMPK signaling pathways and histone H3K9 and H3K14 acetylation (Ac) in male (d) and female (f) mouse hearts and overall lysine acetylation (Ac-K) in male (e) and female (g) mouse hearts. Right panels show quantitative analyses of the expression (n=5 WT, n=6 KI for d-e; n=7 for f-g). Unpaired *t* test. *N* represents biologically independent replicates. *p* values are shown in the figure. Data are mean ± SEM. Source data are provided as a Source Data file.

## Notes

### Competing Interest Statement

The authors have declared no competing interest.

